# Legumain drives processing of cathepsins and nuclear localisation of cathepsin L

**DOI:** 10.1101/2025.08.17.670765

**Authors:** Alexander R. Ziegler, Bangyan Xu, Bethany M. Anderson, Linghui Liu, Stephanie Luedtke, Joanna Sacharz, David A. Stroud, Nichollas E. Scott, Laura E. Edgington-Mitchell

**Author notes:** These authors have contributed equally to this work (Alexander R. Ziegler, Bangyan Xu). Correspondance should be addressed to: Laura Edgington-Mitchell ORCID: 0000-0002-6810-6149.

## Abstract

Lysosomal proteases such as the cathepsin family and the asparaginyl endopeptidase, legumain, govern vital processes to maintain cellular proteostasis, and their dysregulation contributes to diverse pathologies. Recent studies have reported extra-lysosomal localisation of these proteases, especially in the nucleus, cytoplasm, and extracellularly, yet their function is not completely understood. To examine the relationship between legumain and cathepsins, we assessed the activity and expression of cathepsins in wild-type and legumain-deficient (*LGMN^−/−^*) cells using chemical activity-based probes and immunoblots. Processing of cathepsins (CTS) L, V, B, and D from the single-chain to the two-chain form was abrogated in the absence of legumain, with some cell type– and species-specific variation observed. This processing was dependent on legumain activity, although the mechanism remains unclear since recombinant legumain does not appear to directly cleave cathepsins *in vitro*. In cell types where CTSL exists in the nucleus preferentially in its double chain form, loss of legumain led to a reduction in nuclear CTSL levels. To understand the potential role of these lysosomal proteases in the nucleus, we applied our newly refined chemical N-terminomics pipeline, No-enrichment Identification of Cleavage Events (NICE). This analysis revealed widespread changes in both protein abundance and proteolysis, including putative nuclear substrates of CTSL and legumain, that primarily suggest roles in cell proliferation, cell cycle regulation, inflammation, and ribosomal biogenesis. Overall, this study builds on our understanding of the relationship between legumain and cathepsins and provides the first systematic characterisation of lysosomal protease substrates in the nucleus. Our results offer valuable insight into the potential extra-lysosomal roles of these critical proteases.

## Introduction

Lysosomes are a major hub of proteolytic activity^1^, harbouring numerous proteases which are crucial for regulating protein homeostasis and cellular signalling by catalysing proteolysis^2–4^. Among these include the cathepsins (CTS) and an asparaginyl endopeptidase (AEP), legumain^5^. Cathepsins are classified into three families based on their catalytic mechanism; aspartic (D, E), cysteine (B, C, F, H, K, L, O, S, V, W, X/Z), and serine (A, G) cathepsins^6^. Under normal physiological conditions, lysosomal proteases contribute to lysosomal protein turnover^7^, antigen processing, and receptor maturation^8,9^ or cytokine activation associated with immune signalling^10,11^. Dysregulated lysosomal proteolytic activity manifests in a range of pathophysiological conditions such as lysosomal storage disorders^12^ and inflammatory conditions such as autoimmune diseases^13^, neurodegeneration^14^ and cancer^15^. This makes the study of lysosomal proteases crucial for understanding broad cellular functions and highlights their potential as therapeutic targets.

Cathepsins and legumain are typically synthesised as inactive zymogens and trafficked to the endo-lysosomal system in a mannose-6-phosphate (M6P)-dependent manner^16^. Upon entering the acidic milieu of endo-lysosomes, these proteases dissociate from M6P receptors and are activated through removal of their propeptides, often auto-catalytically^17^. Generally, acidic environments are required to maintain the activity of lysosomal proteases, although some cathepsins (e.g., CTSB, CTSS) exhibit proteolytic activity across broader pH ranges^18,19^. Several cathepsins (e.g., B, C, D, H, L, V,) can undergo secondary processing from their single-chain (SC) active form to produce a double-chain (DC) variant consisting of a heavy and light chain that remain associated by a disulfide bond^20^. Both SC and DC forms exhibit proteolytic activity, yet the purpose and impact of this processing step has not been well elucidated^21^. Several studies have demonstrated the dependence of legumain in the processing of CTSB, CTSD, CTSH, and CTSL^7,22–25^. It has long been assumed that legumain directly cleaves the SC to generate the DC, however no direct evidence of this has been reported to date. As such, the interplay between these lysosomal proteases and the potential impact on cellular function remains elusive.

Beyond the lysosome, there is increasing evidence to suggest non-canonical localisation of cathepsins and legumain in the nucleus, cytoplasm, and extracellular milieu^1,26,27^. Most previous research has focussed on extracellular trafficking of lysosomal proteases in the context of cancer^28–30^ and bacterial infection^31,32^, with cytoplasmic or nuclear trafficking and function being less studied. Some cathepsins exhibit alternatively spliced variants lacking the canonical signal peptide, resulting in altered trafficking^33^. CTSB transcript variants lacking the signal peptide localise to the nucleus or cytoplasm^34–36^, and truncated CTSL resulting from alternative AUG codon usage leads to nuclear translocation and processing of CCAAT-displacement protein/cut homeobox transcription factor (CUX1)^37^. Alternatively, nuclear cathepsin localisation can be mediated by an intrinsic nuclear localisation signal (NLS), as with CTSB^35^ or legumain^5^, or by shuttling with NLS-containing proteins such as Snail and BAT3, for CTSL and CTSD, respectively^38,39^. This transport mechanism requires initial cytoplasmic localisation, which may be facilitated by lysosomal membrane permeabilisation (LMP) under stress conditions^40,41^ or by regulated lysosomal leakage^42^.

Several cytoplasmic and nuclear substrates of these proteases have been identified, suggesting their presence and activity within these non-canonical environments. Cytoplasmic CTSB and CTSL have been linked to NLRP3 inflammasome activation^43^. CTSL can cleave dynamin to influence podocyte motility^44^ and legumain is often linked to tau and α-synuclein processing in Alzheimer’s and Parkinson’s disease, respectively^45,46^. Further, legumain, CTSL, and CTSD can process histone H3 in the nucleus, which regulates transcription and cell cycle^47–50^. Nuclear CTSL also drives cancer cell progression by promoting epithelial-to-mesenchymal transition (EMT)^51–53^. Together, emerging reports of extra-lysosomal activity demonstrate the widespread influence of these proteases on cellular function in both normal physiology and disease, although further studies are required to more completely discern their extra-lysosomal roles.

Herein, we evaluate the dependence of legumain on cathepsin processing and the subsequent consequences on their subcellular localisation. While we were unable to observe direct cleavage of cathepsins by legumain *in vitro*, we observed distinct differences in CTSL between wild-type and legumain-deficient cells, whereby loss of legumain impairs CTSL processing to the DC and therefore diminishes its nuclear localisation in some cell types. Using No-enrichment Identification of Cleavage Events (NICE), our newly refined N-terminomics pipeline, we assessed the CTSL– and legumain-specific changes to the proteome and degradome of cytoplasmic– and nuclear-enriched fractions. Our analyses highlight potential functions of these lysosomal proteases in the nucleus, contributing to transcriptional regulation and cell cycle progression through cleavage of nuclear substrates. Together, this study highlights the dependence of legumain protease activity in the processing of various lysosomal cathepsins and the trafficking of CTSL to the nucleus. Our analysis of nuclear-enriched fractions catalogues novel putative substrates of CTSL and legumain in the nucleus, building on our understanding of their extra-lysosomal function.

## Results

### Cathepsin L processing is impaired in the absence of legumain protease activity

To investigate the dependence of legumain on cathepsin processing (Figure 1A), we generated legumain-deficient (*LGMN^−/−^)* HSC-3 human oral cancer cells^54^ (Figure S1A-B, Table S1). We subsequently re-expressed WT (rWT) or catalytically-dead legumain (rC189S, rC-S) in *LGMN^−/−^* HSC-3 cells to assess the requirement for legumain protease activity, as well as a mutant lacking the ability to bind to integrins (rR118H, rHGD). Loss of legumain activity and expression was verified using the legumain-specific activity-based probe (ABP) LE28^55^ and immunoblot, respectively (Figure 1B). Using the pan-cysteine cathepsin ABP BMV109^56,57^, we measured the activity of cathepsins X (CTSX), B (CTSB), L (CTSL), and S (CTSS) (Figure 1C). *LGMN^−/−^* and rC-S HSC-3 cells exhibited a marginal increase in labelling of active cathepsins at approximately 30 kDa compared to cells expressing active legumain (rWT, rHGD) (Figure 1C). Immunoprecipitation revealed that this band contained a mixture of CTSB and CTSL SCs (Figure 1E). BMV109 labelling indicated activity of both the SC and DC CTSL in WT cells, while only the SC was labelled in *LGMN^−/−^* cells. It should be noted here that the catalytic cysteine of CTSB is located within its light chain (∼5 KDa). BMV109 labelling of the CTSB heavy chain is therefore not observed in SDS-PAGE gels even though the DC is expected to be active. Immunoblot revealed a loss of CTSB and CTSL DC and concurrent accumulation of their SC in *LGMN^−/−^* cells (Figure 1C). Cathepsin V (CTSV), a close homologue of CTSL, also remained predominantly in the SC form in the absence of legumain protease activity. A small amount of DC CTSV was observed, suggesting a protease other than legumain may contribute to CTSV processing. rWT and rHGD rescued CTSB, CTSL, and CTSV processing; however, rC-S did not, demonstrating a requirement for active legumain for this processing to occur. On the other hand, cathepsins lacking a DC form in HSC-3 cells (CTSD, CTSX, CTSS) exhibited no alterations to their processing regardless of the presence of legumain (Figure 1D).

**Figure 1.**
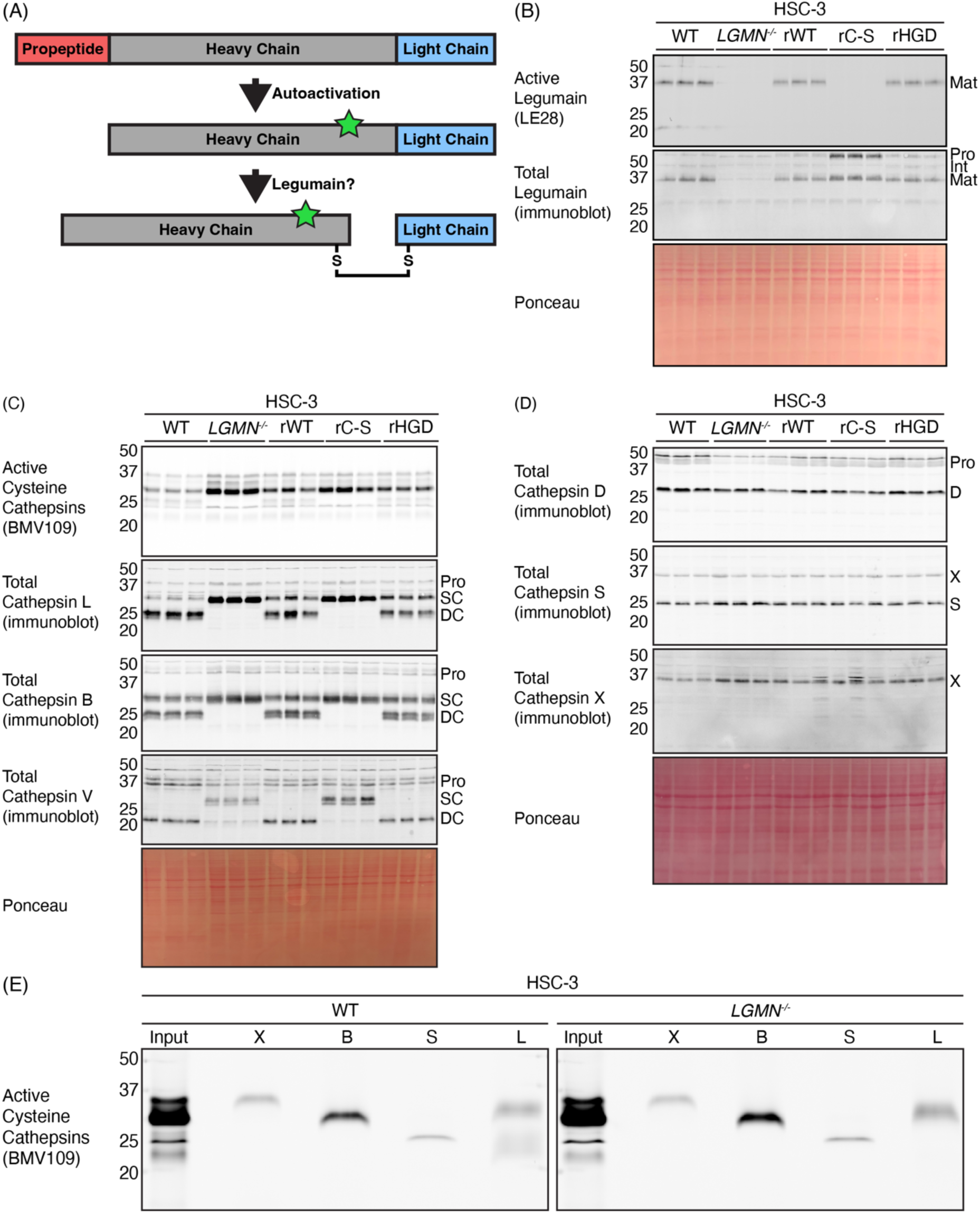
Cathepsin processing is dependent on proteolytically active legumain. (A) Lysosomal proteases such as cathepsins are synthesised as inactive zymogens containing an N-terminal propeptide (red) which inhibits protease activity. Trafficking to the acidic environment of the lysosome promotes auto-activation by removing the propeptide, leading to the generation of an active single chain. This can then be further processed into a double chain variant, which consists of a heavy (grey) and light (blue) chain joined by a disulfide bond (S-S). A schematic of cathepsin L is shown. (B) Lysate labelling of HSC-3 cells with the LE28 activity-based probe to measure legumain activity and immunoblot analysis for legumain expression. Ponceau S stain was used as a loading control (wild-type (WT), legumain-deficient (LGMN^−/−^), and LGMN^−/−^ reconstituted with wild-type legumain (rWT), catalytically-dead mutant (rC189S, rCS) or an integrin-binding mutant (rR118H, rHGD)) (n = 3 per group). (C) Lysate labelling of active cysteine cathepsins in HSC-3 cell constructs using the BMV109 activity-based probe and immunoblot analysis of cathepsins L, B, and V. Ponceau S stain was used as a loading control (n = 3 per group). (D) Immunoblot analysis of cathepsins D, S, and X in HSC-3 cell constructs. Ponceau S stain was used as a loading control. Note that cathepsin S immunoblot was performed on the same membrane following cathepsin X immunoblot (n = 3 per group). (E) Immunoprecipitation of BMV109-labelled WT and LGMN^−/−^ HSC-3 cells with cathepsin X, B, S, and L antibodies. An equal amount of protein was loaded per lane.

Using an additional human oral squamous cell carcinoma line, SCC-9 (Figure S1C), we observed similar processing defects of CTSL, CTSB, and CTSV in the absence of legumain, with no changes to the other cathepsins (D, X, S) (Figure S2). Analysis of spleen tissue from wild-type and *Lgmn^−/−^* mice also demonstrated the dependence of legumain on Ctsb and Ctsl, as well as Ctsd processing (Figure S3A). Immunoprecipitation of these samples revealed increased levels of Ctsb and Ctsl SC (Figure S3B) in *Lgmn^−/−^* mice, consistent with the human oral cancer cells. Genetic deletion of legumain in the murine macrophage cell line, RAW264.7 also resulted in impaired processing of Ctsl and Ctsd (Figure S4). As WT RAW264.7 cells exhibited negligible levels of DC Ctsb, little difference in Ctsb was observed upon legumain deletion (Figure S4B). As was the case for human cells, *Lgmn^−/−^* murine spleen and RAW264.7 cells did not exhibit alterations in Ctsx or Ctss (Figure S3A; Figure S4B). To examine the requirement for catalytically active legumain in murine macrophages, we treated RAW264.7 cells with the legumain-specific inhibitor SD-134^58^. Legumain inhibition led to a reduction in Ctsl and Ctsd DC and accumulation of the SC as observed in the *Lgmn^−/−^* cells (Figure S4C). Overall, these data demonstrate the requirement for proteolytically active legumain to drive the processing of lysosomal cathepsins (B, D, L, V) from the SC to the DC.

### Legumain does not readily cleave cathepsins in vitro

Since the generation of the DC forms of several cathepsins is dependent on legumain protease activity, it has long been hypothesised that legumain directly processes these cathepsins^22,23^. Although the data were not shown, Colbert and colleagues suggested that recombinant legumain could not cleave CTSL SC *in vitro*^59^. To investigate this in more detail, we incubated recombinant CTSL, CTSB, CTSD, or CTSX with activated recombinant legumain in buffers at a range of pH (4.5, 5.5, 6.5) and monitored cleavage by SDS-PAGE (Figure 2A-B; Figure S5A-B). Legumain did not process any of the cathepsins at pH 5.5 following a 5-hour incubation, however, was able to cleave trypsinogen, a previously identified substrate^25^ (Figure 2A). Reducing the pH to 4.5 to more accurately replicate mature lysosomes also failed to promote cathepsin processing (Figure S5A). Additionally, the increase in pH to 6.5 led to complete loss of legumain activity, as demonstrated by the loss of trypsinogen processing (Figure S5B). This supports previous reports of legumain becoming unstable upon reaching environments above pH of 6.0^60^. Denaturing the cathepsins prior to legumain addition led to their degradation, but not double chain generation, suggesting legumain can cleave the unfolded polypeptide chains in these conditions, but not the intact cathepsin proteins (Figure 2B). We also added recombinant legumain (rLGMN) to acidified protein lysates (citrate buffer, pH 5.5) collected from WT, *LGMN^−/−^*, rWT, rC-S and rHGD HSC-3 cells, which would more faithfully recapitulate the legumain-cathepsin interaction, including potential cofactors or post-translational modifications that may be required for cathepsin processing (Figure S5C). Endogenously cleaved DC CTSL was only observed in lysates from cells expressing active legumain, and addition of rLGMN did not provoke further CTSL processing in any cell line. It is clear from the Ponceau stain that legumain addition led to cleavage/degradation of many other proteins under these conditions, and in fact, pro-CTSL appears to be trimmed upon addition of legumain. Since our *in vitro* assays were unable to recapitulate legumain-dependent generation of DCs, we hypothesised that native cellular conditions may be required for this processing to occur.

**Figure 2.**
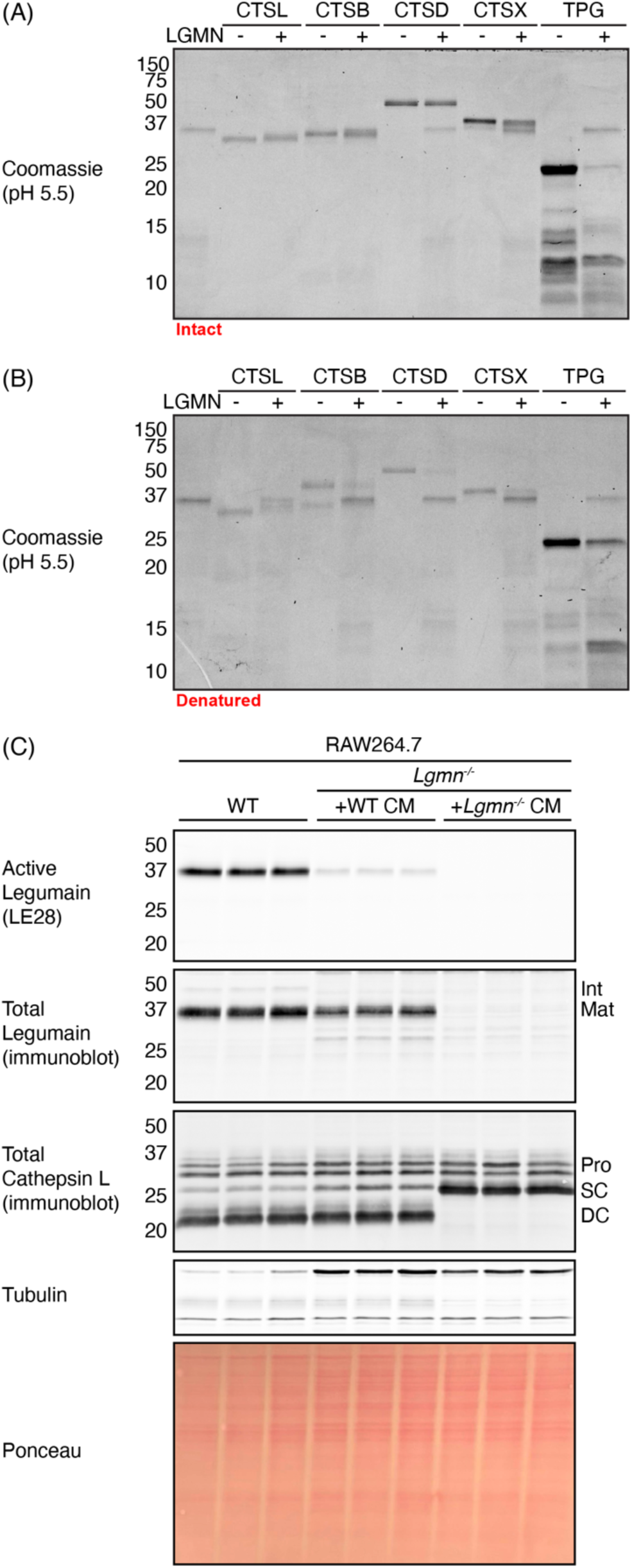
Legumain does not directly cleave lysosomal cathepsins in vitro. (A) In vitro cleavage assay of intact proteins using activated recombinant legumain (LGMN) and recombinant cathepsins L (CTSL), B (CTSB), D (CTSD), and X (CTSX) at pH 5.5. Trypsinogen (Tpg) was used as a positive control (n = 3 independent experiments). (B) In vitro cleavage assay of denatured proteins at pH 5.5. Proteins were boiled at 95°C for 5 minutes prior to addition of legumain and incubation. Gels were visualised by Coomassie stain. (C) Assessment of legumain uptake and activity, and its ability to process cathepsin L in legumain-deficient (Lgmn^−/−^) RAW264.7 cells. Activity was measured using the activity-based probe LE28, followed by immunoblot of legumain and cathepsin L to assess expression. Tubulin and Ponceau S stain were used as loading controls (n = 3 per group).

To investigate the dependence on native cellular environments, we collected conditioned media (CM) from WT and *Lgmn^−/−^* RAW264.7 cells, which contains legumain secreted in the inactive zymogen form (Figure S6A). We applied this media to *Lgmn^−/−^* RAW264.7 cells for 48 hours to allow legumain uptake before analysing CTSL processing. WT CM, but not *Lgmn^−/−^* CM, led to clear uptake and activation of legumain, though less than in untreated WT cells (Figure 2C). Even this low level of active legumain led to nearly complete rescue of Ctsl processing to the DC form. We also observed this rescue in HSC-3 cells, albeit to a much lesser extent than RAW264.7 cells (Figure S6B-D). This likely reflects the enhanced ability of macrophages to engulf exogenous material compared to epithelial cells^61,62^. These data confirm that active legumain in the endolysosomal compartment can drive cathepsin processing; however, the interaction may require as yet unknown cofactors, specific physicochemical conditions, or intermediary proteases.

### Cathepsin L is preferentially trafficked to the nucleus in the double chain form

We recently observed that CTSL exists in the nucleus predominantly in its DC form in Mutu murine dendritic cells^63^. Since Mutu cells lacking Ctsx exhibited less DC Ctsl and less nuclear Ctsl, we hypothesised that *LGMN^−/−^* cells would exhibit a similar phenotype. Immunoblotting revealed changes in the proportion of CTSL SC and DC levels between cytoplasmic and nuclear-enriched fractions, with nuclear CTSL almost completely absent in nuclear fractions of *LGMN^−/−^* cells (Figure 3). This appeared to be selective for CTSL, since total nuclear levels of CTSB and CTSV were not impacted by loss of legumain. These results were reproducible in SCC-9 cells with the DC being the primary form of CTSL present in nuclear-enriched fractions in WT cells, and minimal nuclear CTSL observed in *LGMN^−/−^* cells (Figure S7). The presence of other lysosomal cathepsins such as CTSD and CTSX in nuclear-enriched fractions was minimal. CTSL was also observed predominately in the DC form in nuclear-enriched fractions collected from WT RAW264.7 cells. In contrast to HSC-3 and SCC-9 cells, however, nuclear Ctsl SC was readily detected in *Lgmn^−/−^* RAW264.7 cells (Figure S8A). Overall, these data suggest that CTSL may preferentially be trafficked to the nucleus in its DC form in a cell-specific manner, and the legumain-dependent processing of CTSL may impact its nuclear localisation and therefore function in these cells.

**Figure 3.**
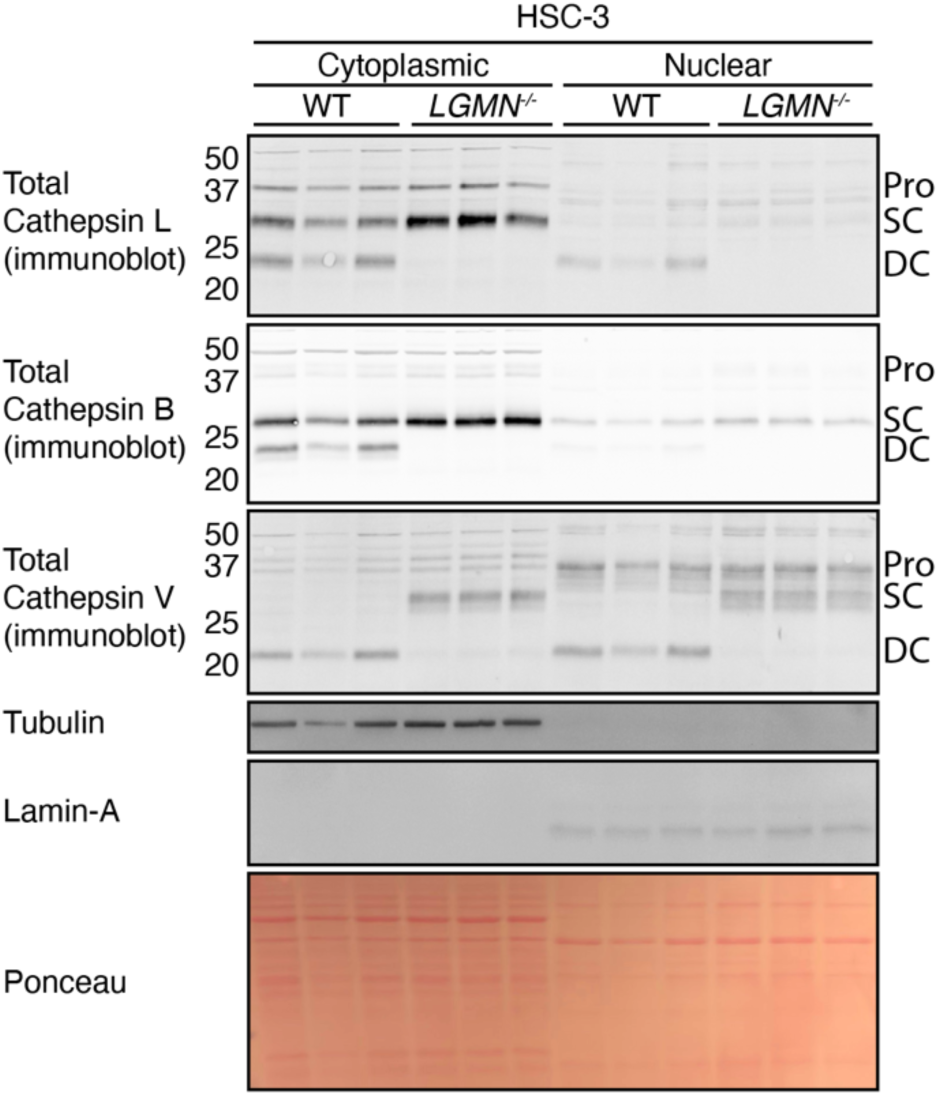
Cathepsin L trafficking to the nucleus is altered in the absence of legumain. Following subcellular fractionation of wild-type (WT) and legumain-deficient (LGMN^−/−^) HSC-3 cell lysates, immunoblot analysis of cathepsins L, B, and V was performed to assess their total levels. Tubulin and lamin-A were used as markers for cytoplasmic and nuclear fractions, respectively. Ponceau S stain was used as a loading control (n = 3 per group).

To examine the trafficking route of nuclear Ctsl, we treated nuclear-enriched and cytoplasmic-enriched RAW264.7 fractions with the glycosidase PNGase F, which removes N-linked glycan modifications. Ctsl in both fractions exhibited an identical shift in gel mobility, suggesting both cytoplasmic and nuclear Ctsl are glycosylated (Fig. S8B). This indicates that nuclear Ctsl traffics through the Golgi where it is targeted to endolysosomes, rather than being synthesised directly in the cytoplasm. While the mechanism remains unclear, it is possible that lysosomal membrane permabilisation (LMP) contributes to release of Ctsl into the cytoplasm for nuclear import by hitchhiking with Snail or other NLS-containing proteins^38^.

### Loss of cathepsin L in oral cancer cells impacts the nuclear proteome

Since the presence of nuclear CTSL was altered in *LGMN^−/−^*cells (Figure 3), we sought to investigate the function of these lysosomal proteases in the nucleus by systematically characterising their subcellular impact on protein abundance and cleavage events. Considering numerous studies have implicated nuclear CTSL function^38,48,64^, along with our observation that CTSL trafficking is distinct from other cathepsins, we first aimed to examine the impact of this protease. To evaluate the cellular changes mediated by CTSL, we generated CTSL knockout HSC-3 cells using the CRISPR/Cas9 system^54^ (Figure S9A, Table S1). *CTSL^−/−^*cells showed a reduction in the bands corresponding to CTSL following live labelling with BMV109 (Figure S9B). Immunoblot analysis also confirmed loss of CTSL expression in these cells. Concurrently, lysate labelling with BMV109 revealed upregulated activity of cathepsins B, S, and X in the absence of CTSL (Figure S10A). CTSL is not reliably labelled by ABPs in cell lysates, most likely indicating its instability following cell lysis. The increased activity of CTSB and CTSS corresponded to an increase in their total expression levels (Figure S10A). Using a CTSX-selective ABP, sCy5-Nle-SY^65^, we confirmed the increase in CTSX activity, and this corresponded to increased CTSX expression (Figure S10B). Likewise, the activities of cathepsin C (CTSC) and legumain were increased in *CTSL^−/−^*cells (Figure S10C-D), as well as total levels of the mature form of legumain (Figure S10D). Together, these data indicate upregulated lysosomal protease expression in the absence of CTSL, most likely as a compensatory mechanism.

We recently developed an enrichment-free N-terminomics workflow to simultaneously assess protein abundance changes and detect proteolytic cleavage events using a chemical tag for proteolysis^66^. Previously, the use of fractionation methods such as high-field asymmetric waveform ion mobility spectrometry (FAIMS) or basic reverse-phase fractionation were used to facilitate increased proteome coverage, thereby achieving sufficient depth to unveil more protease cleavage events as compared to unfractionated samples^66,67^. Herein, we capitalise on recent technological advancements introduced in the Orbitrap Astral^68^, which is vastly faster and more sensitive, to circumvent the need for both enrichment and fractionation. Use of data-independent acquisition (DIA) permitted deep proteome and N-terminome coverage in a fraction of the time (70 minutes compared to 12 hours per sample), using a fraction of the input (200 ng compared to 12 µg). We henceforth refer to this method as No-enrichment Identification of Cleavage Events (NICE) (Figure 4).

**Figure 4.**
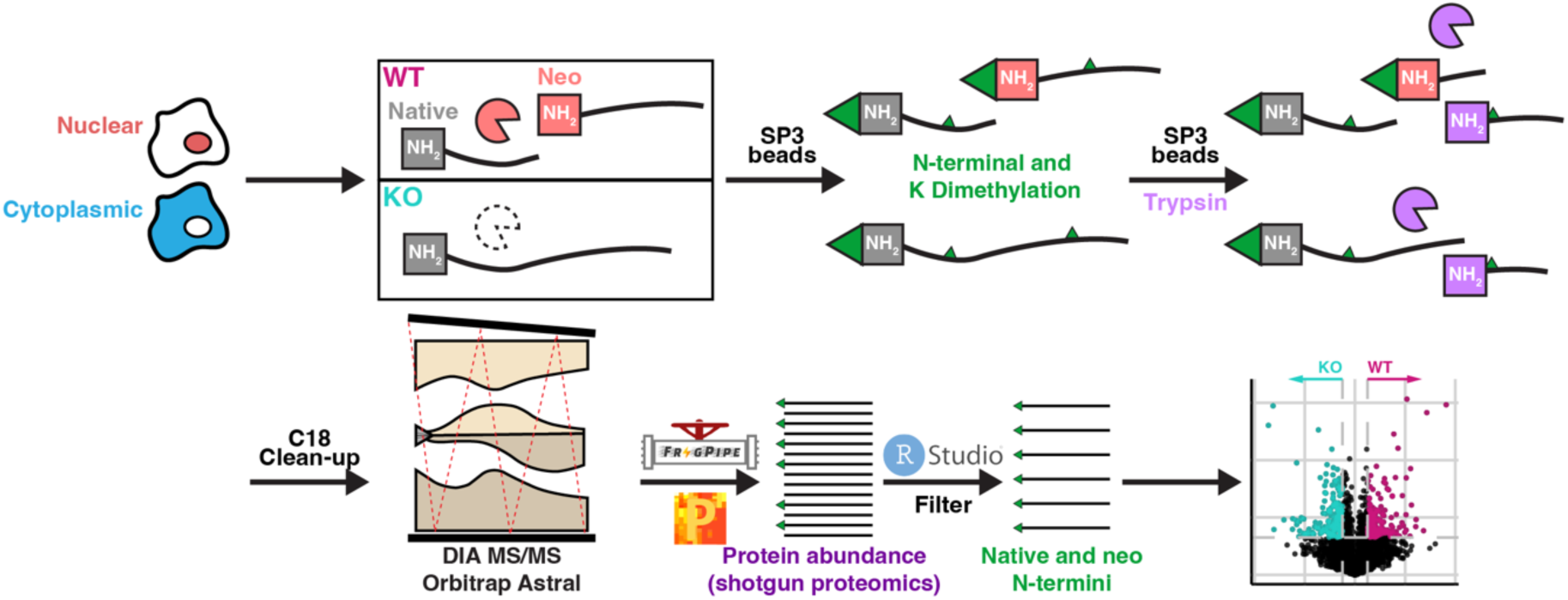
No-enrichment Identification of Cleavage Events (NICE) analysis to assess protein abundance changes and cleavage events. Schematic of the NICE workflow. Following subcellular fractionation, lysates were dimethylated at the protein level (green) to label native and protease-generated (neo) N-termini, as well as lysine (K) side chains. In vitro trypsin digestion and de-salting with C_18_ was performed for analysis by mass spectrometry using data-independent acquisition (DIA) on the Orbitrap Astral. Data analysis was performed using FragPipe, Perseus, and RStudio. All peptides were analysed to quantify protein abundance, while those with N-terminal dimethylation were filtered for cleavage site analysis.

Following subcellular fractionation of WT and *CTSL^−/−^*HSC-3 cells, lysates were subjected to NICE analysis, identifying 4,897 proteins from 38,994 peptides in nuclear-enriched fractions and 5,718 proteins from 49,573 peptides in cytoplasmic-enriched fractions (Figure 5A). These data revealed significant changes in protein abundance within each fraction (Figure 5B-C; Table S2-3). Many of the proteins were identified in both subcellular fractions with 1,486 unique to the nuclear-enriched fractions and 2,307 solely identified in cytoplasmic-enriched fractions (Figure 5D). *CTSL^−/−^* HSC-3 cells exhibited a significant increase of CTSS and CTSB in both fractions, as well as trending increases in other lysosomal cathepsins, validating the immunoblot analysis on whole-cell lysates (Figure S10). Of the *CTSL^−/−^*-enriched proteins, one of the most highly enriched proteins in both fractions was guanylate-binding protein 1 (GBP1), which is primarily involved in inflammasome and autophagy activation in response to foreign pathogens^69^ (Figure 5B-C).

**Figure 5.**
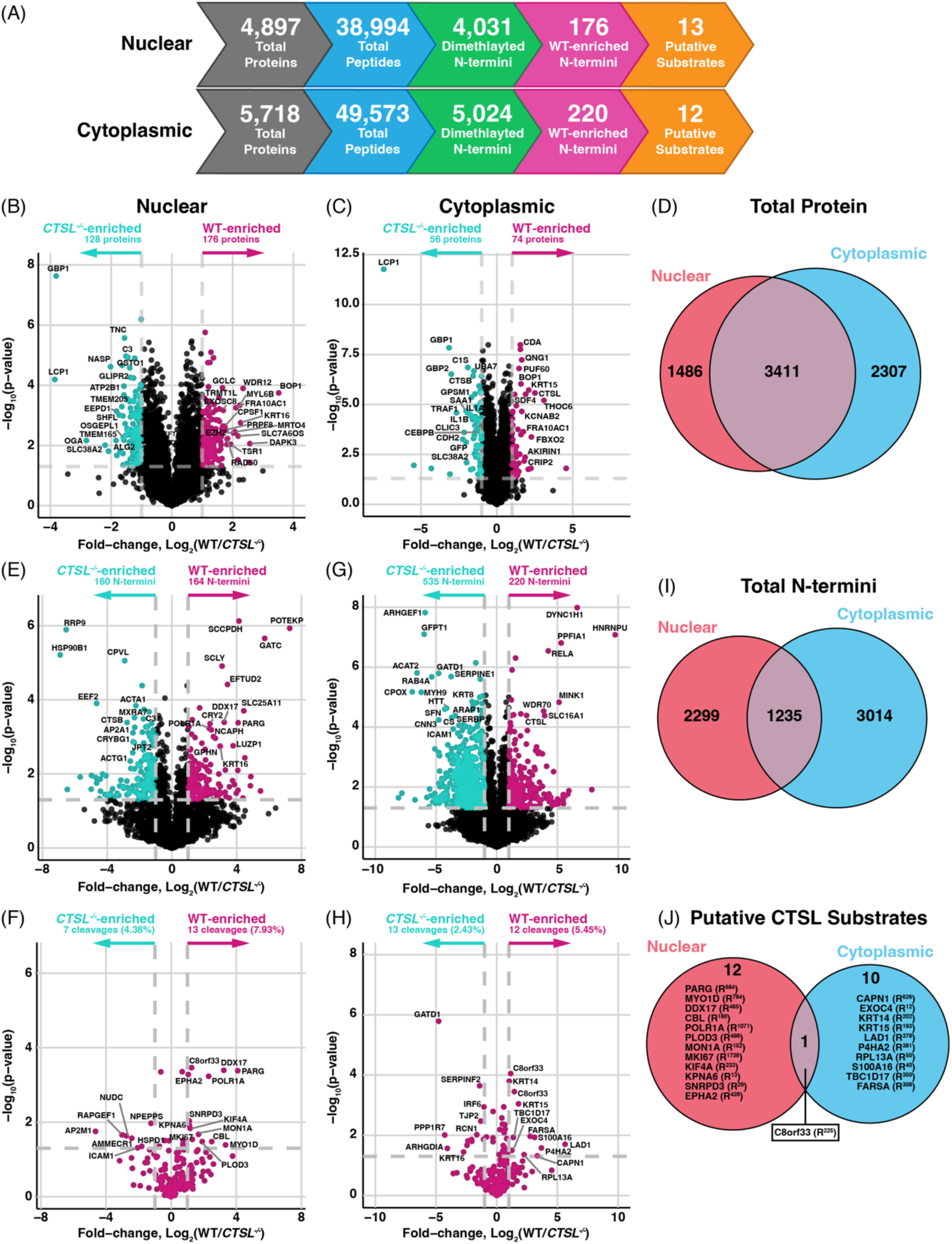
Protein abundance and proteolytic changes between wild-type (WT) and cathepsin L-deficient (CTSL^−/−^) HSC-3 cells suggest potential roles of cathepsin L in the nucleus. (A) Total protein and peptide identifications per subcellular fraction were summarised. Peptides were filtered for dimethylation at the N-terminus, indicating native and neo-N-termini. (B-C) Volcano plots of protein abundance differences between WT and CTSL^−/−^ nuclear-enriched (B) and cytoplasmic-enriched (C) fractions (n = 4 per group). A Student’s two-way t-test was used to determine significance at |Log_2_(WT/CTSL^−/−^) > 1| and –log_10_(p-value) 1.3 (p < 0.05). (D) Overlap of total proteins identified between nuclear– and cytoplasmic-enriched fractions. (E-H) Peptides identified by No-enrichment Identification of Cleavage Events (NICE) were filtered for those dimethylated at the N-terminus, representing native and protease-generated (neo) N-termini (n = 4 per group). (E, G) Volcano plots of N-termini differences between WT and CTSL^−/−^ nuclear-enriched (E) and cytoplasmic-enriched (G) fractions. A Student’s two-way t-test was used to determine significance at |Log_2_(WT/CTSL^−/−^) 1| and –log_10_(p-value) > 1.3 (p < 0.05). (F, H) N-termini which result from potential cathepsin L cleavage, according to the hypothesised cathepsin L cleavage motif (XX(F/W/Y)(K/R)τXXXX) are highlighted in magenta for nuclear-enriched (F) and cytoplasmic-enriched (H) fractions. (I) Overlap of total N-termini detected in nuclear– and cytoplasmic-enriched fractions from WT and CTSL^−/−^ HSC-3 cells. (J) Overlap of putative cathepsin L (CTSL) substrates identified in nuclear– and cytoplasmic-enriched fractions.

Among the 176 WT-enriched proteins in the nuclear-enriched fractions, many were proteins corresponding to ribosomal subunits, such as RPL22, RPL23A, RPS6, RPS8, RPS1, RPS24, and others (Figure 5B; Table S2). The most significantly increased protein in WT nuclear-enriched fractions was ribosome biogenesis protein BOP1 (BOP1), which functions to form and mature these ribosomal subunits in the nucleus^70^. Transforming growth factor-beta (TGF-β) and a range of motor proteins such as myosin and kinesin chains (MYL6, MYL6B, MYL12B, MYH9, KLC1, and KLC2) were also upregulated in WT nuclear-enriched fractions. Together, these protein abundance changes offer insight into how CTSL contributes to TGF-β signalling or increased cell migration, which promote cancer progression. As such, these data will be useful in discerning CTSL-specific effects and mark the first proteome catalogue for the interrogation of the role of CTSL in oral cancer cells.

### Cleavage events within nuclear-enriched fractions reveal putative roles for cathepsin L in the nucleus

To identify potential nuclear substrates of CTSL, we assessed the N-terminome differences of nuclear-enriched fractions from WT and *CTSL^−/−^* HSC-3 cells following NICE analysis (Figure 5E; Table S4). As peptides are dimethylated at the protein-level in the NICE workflow (Figure 4), this provides a chemical tag for proteolysis to identify protease-mediated cleavages events. Of the 38,994 peptides detected across the replicates, 4,031 were dimethylated at the N-terminus (10.34%), indicating native or neo-N-termini (Figure 5A; Figure S11A). We observed dimethylation efficiency of >92% in each sample (Figure S11B). Of the detected N-termini, 164 were enriched in WT, indicating cleavage events facilitated directly by CTSL or other proteases that it regulates (Figure 5E). The majority of these N-termini resulted from cleavage following arginine residues, likely representing cathepsin or trypsin-like protease activity (Figure S11C). A similar cleavage motif was also observed among the 160 *CTSL^−/−^*-enriched N-termini (Figure S11D). GO and pathway analyses revealed these WT-enriched N-termini were broadly associated with mRNA splicing and protein translation (Figure S11E-F), whilst the *CTSL^−/−^*-enriched N-termini exhibited enrichment of translation and trafficking mechanisms (Figure S11G-H).

In attempt to identify direct CTSL substrates in the nucleus, we filtered for N-termini based on the predicted CTSL cleavage motif, XX(F/W/Y)(K/R)τXXXX^71^. We identified 13 putative CTSL substrates enriched in nuclear-enriched WT fractions (Fig. 5F; Table S4). These included cleavage of proteins related to mRNA/RNA splicing such as SNRPD3, DDX17, and POLR1A, and proteins associated with chromosomes or chromatin during mitosis and cell proliferation such as proliferation marker protein Ki-67 (MKI67) and chromosome-associated kinesin KIF4A. We also observed cleavage of the nuclear import protein importin subunit alpha-7 (KPNA6) at R^13^τM^14^. These proteins may represent putative novel substrates of CTSL in the nucleus, however since CTSL has a broad substrate specificity, there may also be additional substrates identified within our WT-enriched N-termini which are cleaved outside this motif.

In the cytoplasmic-enriched fractions of the same cells, we observed significant upregulation of CTSB, CTSC, and CTSS in the absence of CTSL, whereas CTSL was enriched in WT cells as expected (Figure 5C). This led to many altered N-termini in the presence and absence of CTSL (Figure 5G; Table S5), which were distinct from those identified in the nuclear-enriched fractions (Figure 5I). The dimethylation efficiency was >92% (Figure S12A). Of the altered N-termini, transcription factor p65 (RELA) isoform 3 at R^294^τT^295^ was enriched in WT samples (Figure 5G), which may impact nuclear factor kappa B (NFκB) signalling. Like the nuclear-enriched fractions, the cleavage motifs were dominated by arginine in the P1 position, however there were minor shifts for preference of phenylalanine and leucine in P1, valine in P2, and asparagine in P3 in the *CTSL^−/−^* cells (Figure S12B-C). GO and pathway analysis revealed enrichment of ribosomal function or nucleotide metabolism for WT-enriched N-termini, suggesting CTSL may contribute to these pathways (Figure S12D-E).

In the absence of CTSL, we primarily observed inflammatory and immune-related responses as seen in the protein-level data, which may be a result of the compensatory upregulation of other lysosomal proteases (Figure S12F-G). We filtered for putative CTSL substrates and observed a greater proportion of these in the WT lysates (Figure 5H). Twelve putative CTSL substrates were enriched in WT fractions, one of which was also observed in the nuclear-enriched fractions, cleavage of isoform 2 of UPF0488 protein C8orf33 (C8orf33) at R^225^τA^226^ (Figure 5J). We also detected the cleavage of calpain-1 (CAPN1) at R^625^τK^626^ as a putative CTSL substrate in the cytoplasmic-enriched fractions (Figure 5H), which may have cascading proteolytic effects downstream of CTSL. Overall, our data suggest diverse roles for CTSL in the nucleus and cytoplasm, providing a basis for further investigation and validation of its proteolytic contribution in these extra-lysosomal compartments.

### NICE analysis reveals legumain-mediated proteolytic events in oral cancer cells

As legumain influences nuclear CTSL (Figure 3), we reasoned that the nuclear substrate repertoire of CTSL would be altered between WT and legumain-deficient (*LGMN^−/−^*) HSC-3 cells. We first confirmed the presence and activity of legumain in HSC-3 cell nuclei by live labelling with LE28 (Figure S13A). In attempt to identify proteolytic events which may result from CTSL in this context, we applied NICE to nuclear– and cytoplasmic-enriched fractions collected from WT and *LGMN^−/−^* HSC-3 cells. We identified 3,125 proteins from 21,505 peptides in the nuclear-enriched fractions (Figure 6A), with 186 upregulated in the WT cells and 98 in the *LGMN^−/−^* cells (Figure 6B; Table S6). Many of the WT-enriched proteins were splicing factors such as SF3B1, U2AF2, SRSF11, SFSWAP, SF3B2, and SF3B3, as well as those involved in chromatin organisation including SMARCA4, SMARCB1, SMARCC1, and SMARCE1. These data suggest legumain may regulate transcription and mRNA splicing by altering the abundance of these nuclear proteins. In the cytoplasmic-enriched fractions, we detected 4,275 proteins from 32,433 peptides (Figure 6A), including 55 WT-enriched and 70 *LGMN^−/−^*-enriched proteins (Figure 6C; Table S7). Many proteins associated with microtubule function were enriched in the presence of legumain including MAP2, KIF2C, and the tubulin subunit TUBB2B. Conversely, we observed proteins relating to immune function or proteolysis in the *LGMN^−/−^* cytoplasmic-enriched fractions, including lysosomal proteins CTSS, progranulin (PRG), acid ceramidase (ASAH1), and sphingomyelin phosphodiesterase (SMPD1) (Figure 6B). Like the previous comparison with *CTSL^−/−^*, many of the identified proteins overlapped between nuclear– and cytoplasmic-enriched fractions (Figure 6D). Together, our data implicate the upregulation of lysosomal proteins in the absence of legumain to restore lysosomal homeostasis, whilst cytoplasmic legumain may promote changes to the cell cytoskeleton.

**Figure 6.**
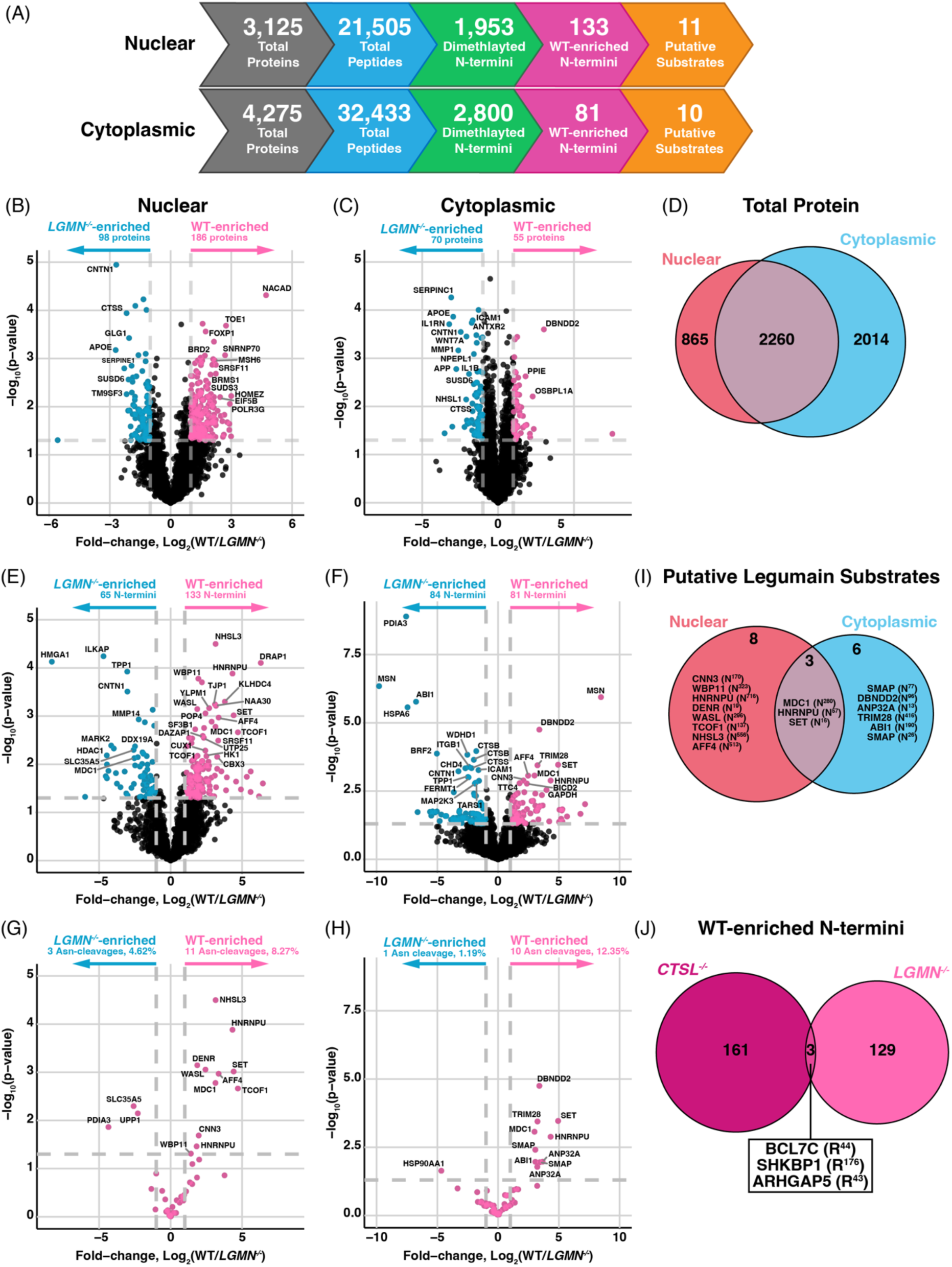
No-enrichment Identification of Cleavage Events (NICE) analysis of wild-type (WT) and legumain-deficient (LGMN^−/−^) HSC-3 cells reveals proteolytic contributions of legumain in the nucleus. (A) Total protein and peptide identifications per subcellular fraction were summarised. Peptides were filtered for dimethylation at the N-terminus, indicating native and neo-N-termini. (B-C) Volcano plots of protein abundance differences between WT and LGMN^−/−^ nuclear-enriched (B) and cytoplasmic-enriched (C) fractions (n = 4 per group). A Student’s two-way t-test was used to determine significance at |Log_2_(WT/LGMN^−/−^) > 1| and – log_10_(p-value) > 1.3 (p < 0.05). (D) Overlap of total proteins identified between nuclear– and cytoplasmic-enriched fractions. (E-H) Peptides identified by No-enrichment Identification of Cleavage Events (NICE) were filtered for those dimethylated at the N-terminus, representing native and protease-generated (neo) N-termini (n = 4 per group). (E-DF) Volcano plots of N-termini detected in nuclear-enriched (E) and cytoplasmic-enriched (F) HSC-3 lysates. A Student’s two-way t-test was used to determine significance at |Log_2_(WT/LGMN^−/−^) > 1| and – log_10_(p-value) > 1.3 (p < 0.05). (G-H) N-termini arising from cleavage following asparagine are highlighted in pink for nuclear-enriched (G) and cytoplasmic-enriched (H) fractions, representing putative legumain substrates. (I) Overlap of putative legumain substrates identified in cytoplasmic-enriched and nuclear-enriched fractions from HSC-3 cells. (J) Overlap of nuclear WT-enriched N-termini in the comparison between WT/CTSL^−/−^ and WT/LGMN^−/−^ analyses, indicating proteins which may be directly processed by cathepsin L in the nucleus.

To gain greater insight on the proteolytic role of legumain in these subcellular compartments, we assessed their respective N-terminomes. We first focussed on N-terminomic analysis of the nuclear-enriched fractions to identify cleavage events facilitated directly or indirectly by legumain, with the indirect events potentially arising due to the differences in CTSL levels between the two genotypes. The dimethylation efficiency was observed to be >90% these fractions (Figure S13B). Of the 21,505 peptides detected in the nuclear-enriched fractions, 1,953 (9.08%) were dimethylated at their N-terminus, indicating native or protease-generated (neo) N-termini (Figure 6A; Figure S14A). Our statistical comparison highlighted 133 N-termini enriched in the presence of legumain, which would indicate cleavage events directly mediated by legumain or indirectly through the proteases that it regulates (Fig. 6E; Table S8). By comparison, 65 N-termini were increased in *LGMN^−/−^* nuclear-enriched fractions. In line with the unique asparaginyl endopeptidase activity of legumain^5^, we observed a clear enrichment in N-termini resulting from cleavage after asparagine in the WT fractions (Figure S14B). This was also observed in the sequence logos, with a marginal preference for asparagine at the P1 position in the WT-enriched, but not the *LGMN^−/−^*-enriched N-termini (Figure S14C-D). Gene ontology (GO) and pathway analyses revealed processes corresponding to RNA splicing regulation and the spliceosome complex in the presence of legumain (Figure S14E-F), whereas wound healing, coagulation, haemostasis, and lysosomes were associated with the *LGMN^−/−^* cells (Figure S14G-H), similar to the *CTSL^−/−^* cells (Figure S11G-H).

In the cytoplasmic-enriched fractions, we observed >90% efficiency in dimethyl labelling (Figure S15A), leading to the detection of 81 N-termini enriched in the WT cells and 84 enriched in *LGMN^−/−^* cells (Figure 6F; Table S9). Like the nuclear-enriched fractions, there was a greater preference for asparagine at the P1 position in the WT-enriched N-termini (Figure S15B-C), representing legumain-mediated proteolytic events. Likewise, the biological processes and pathways associated with the WT-enriched and *LGMN^−/−^*-enriched N-termini were similar to the nuclear-enriched fractions (Figure S15D-G).

To identify which of these cleavage events could be directly mediated by legumain, we filtered for N-termini arising from cleavage after asparagine residues, reflecting its asparaginyl endopeptidase activity (Figure 6G-H). We identified 11 asparaginyl cleavages enriched in the WT nuclear-enriched fractions whilst only 3 were detected in the *LGMN^−/−^* (Figure 6G). Additionally, ten putative legumain substrates were identified in the cytoplasmic-enriched fractions (Figure 6H). Of the putative legumain substrates, we observed cleavage of heterogenous nuclear ribonucleoprotein U (HNRNPU) at N^57^τG^58^, which we have previously identified as a substrate in the murine spleen^66^. We also identified an additional putative legumain substrate in both nuclear– and cytoplasmic-enriched fractions, mediator of DNA damage checkpoint protein 1 (MDC1), which was cleaved at N^280^τG^281^ and is a protein known to contribute to DNA damage repair^72^ (Figure 6I).

Since nuclear CTSL is reduced in *LGMN^−/−^* oral cancer cells (Figure 3), we reasoned that N-termini enriched in WT cells compared to both *LGMN^−/−^* and *CTSL^−/−^* cells may provide greater confidence in identifying nuclear CTSL substrates. Only 3 cleavage events were observed to be lost in both *LGMN^−/−^* and *CTSL^−/−^*cells (Figure 6J). These included processing of B-cell CLL/lymphoma 7 protein family member C (BCL7C) at R^44^τI^45^, SH3KBP1-binding protein 1 (SHKBP1) at R^176^τM^177^, and Rho GTPase-activating protein 5 (ARHGAP5) at R^43^τS^44^. Of these, the protein abundance of BCL7C and ARHGAP5 follows a similar trend to the observed N-termini (Table S6), indicating the increased cleavages observed in these cells may be a result of increased protein expression rather than differential proteolysis. Similar trends were also observed in the comparison between WT and *CTSL^−/−^* nuclear-enriched fractions (Table S2). We also searched for known CTSL nuclear substrates in our N-terminomics data and found cleavage of CUX1 at R^1112^τV^1113^ in isoform 2 to be significantly upregulated in nuclear-enriched fractions from WT compared to *LGMN^−/−^* cells (Figure 6E; Table S8). This peptide was not detected in the WT/*CTSL^−/−^* comparison (Table S4). This detected cleavage site is distinct from those previously identified^37^ at Q^643^τE^644^, S^747^τT^748^, and S^755^τS^756^, and therefore may offer an additional CTSL-mediated site or CTSL-independent proteolytic event to regulate CUX1 function. These data collectively highlight global changes to the proteome and N-terminome of nuclear– and cytoplasmic-enriched fractions in oral cancer cells dependent on the lysosomal proteases legumain and CTSL. The direct cleavage of these proteins, and their dependence on the CTSL DC variant specifically warrants further investigation to understand the role of CTSL in the nucleus.

## Discussion

Our study aimed to elucidate the impact of legumain protease activity on the processing of various lysosomal cathepsin proteases, the consequential effect on CTSL trafficking to the nucleus, and its subsequent roles there. We observed a requirement for legumain activity in processing CTSL, CTSB, and CTSV from the SC to the DC in human oral cancer cell lines, and murine Ctsd was also affected (Figure 1; Figure S4). Our *in vitro* cleavage assays revealed that legumain does not directly cleave CTSL, CTSB, and CTSD, at least in isolation (Figure 2A). Addition of exogenous pro-legumain to legumain-deficient cells, however, rescued processing of CTSL (Figure 2C; Figure S6).

We previously observed that Ctsl SC accumulated in Ctsx-deficient murine dendritic cells, similar to the impact of legumain loss but to a lesser extent, and this was recapitulated by Ctsx inhibitors^63^. Loss of Ctsx activity had no measurable impact on legumain activity and Ctsx cannot directly process Ctsl due to its status as a carboxy-exopeptidase^63^. Together these data suggest that additional cofactors or precise cellular environments may be required for SC-to-DC conversion to occur. Alternatively, there may be intermediary proteases regulated by legumain that mediate the processing. Similar to previous studies^7,73,74^, we observed widespread upregulation of lysosomal proteases in *LGMN^−/−^* HSC-3 cells, most likely as a compensatory response to maintain lysosomal homeostasis. As such, the altered lysosomal environment resulting from LGMN or CTSX deletion may impair CTSL processing. The complex interplay between these lysosomal proteases and the consequences of CTSL processing requires further investigation to more completely understand lysosomal biology.

While we could not determine the definitive mechanism for legumain-dependent processing of CTSL, we next aimed to explore the consequences of this relationship. In murine dendritic cells, we previously observed that Ctsl exists in the nucleus predominately in the DC form, and that loss of Ctsx activity resulted in less DC formation and therefore less nuclear Ctsl. We hypothesised that loss of legumain activity would similarly impact nuclear CTSL levels. Indeed, in two independent oral cancer cell lines, nuclear CTSL was exclusively in its DC form, and we observed loss of nuclear CTSL in the absence of legumain (Figure 3; Figure S7). In murine RAW264.7 macrophages, however, Ctsl was present in the nucleus in pro, SC, and DC forms Figure S8). Here, legumain deficiency did not impact the total level of nuclear Ctsl, only its processing, which indicates cell-type-specific differences in nuclear CTSL trafficking. Binding to the NLS-containing protein Snail is thought to permit importin-dependent nuclear transport of Ctsl^38^. It is possible that this mechanism is more favoured in the DC form, and that mechanisms may differ across cell types.

To gain further insight into the relationship between CTSL and legumain, we applied our refined N-terminomics pipeline, NICE, to cytoplasmic– and nuclear-enriched fractions. We hypothesised that processing of nuclear CTSL substrates would be impaired in *LGMN^−/−^* cells due to the reduction of nuclear CTSL in these cells. Unexpectedly, we observed only three overlapping cleavage sites enriched in the WT nuclear-enriched fractions (Figure 6J). The minimal overlap in these cleavage sites could suggest several possibilities: the expected CTSL DC substrates could be cleaved outside the nucleus prior to import, meaning the dependence of legumain is negligible; the resulting proteolytic products could undergo nuclear export following processing and therefore are undetectable in the nuclear-enriched fractions; or there may be redundancy in CTSL-mediated cleavage events, potentially with other cysteine cathepsins such as CTSB, with which CTSL shares similar cleavage specificities and functions^75^. One of these overlapping nuclear substrates, BCL7C, harbours anti-apoptotic properties, whereby its downregulation in ovarian cancer facilitates worse prognoses for patients due to increased cellular proliferation and invasion^76^. BCL7C is also downregulated in other cancer types, and as such, it is a recognised tumour suppressor^77^. Degradation of this protein may prove detrimental and contribute to differences in the proliferative ability of the cells. Likewise, the detected cleavage of SHKBP1 may also contribute to these processes through its ability to promote epithelial growth factor receptor (EGFR) signalling and cellular differentiation^78,79^. Further cleavage validation of these putative substrates by CTSL using *in vitro* assays are required to confirm direct proteolysis.

Despite observing less-than-anticipated overlap in the cleavage events lost in *LGMN^−/−^* and *CTSL^−/−^* cells compared to WT, the rich datasets provide valuable insight in deciphering the role and cell biology of both CTSL and legumain. Many of the protein abundance changes between WT and *CTSL^−/−^* cells suggest roles for CTSL in inflammation and cell cycle regulation. GBP1 was one of the most highly upregulated proteins in *CTSL^−/−^*nuclear-enriched and cytoplasmic-enriched fractions (Figure 5B-C). Elevated GBP1 expression is recognised as a biomarker in oral cancer^80,81^, linking to worse prognosis^82^. GBP1 is recruited to lysosomes following endo-lysosomal damage^83^ and given the changes to lysosomal homeostasis in *CTSL^−/−^* cells, it may facilitate GBP1 upregulation. Previously, GBP1 has been linked to CTSB whereby inhibiting CTSB downregulated GBP1 and pro-inflammatory protein expression^84^. Conversely, we observed increased CTSB in *CTSL^−/−^* cells (Figure S10A), which may facilitate upregulation of GBP1 and the pro-inflammatory proteins IL-1β, IL-1α, CEBPB, ICAM1, and C3 in the cytoplasmic-enriched fractions (Figure 5C; Table S3).

To further support the role of CTSL in inflammation, we observed increased processing of RELA/p65 in WT cytoplasmic-enriched fractions (Figure 5G). RELA/p65 is part of the NFκB complex which transcriptionally regulates various pro-inflammatory genes. The observed cleavage at R^294^τT^295^ matches R^304^τT^305^ in the canonical isoform and occurs C-terminal to the NLS. This may therefore impact the nuclear translocation of RELA/p65 and reduce its transcriptional activation, as seen with N-terminal processing by copine-I^85^. Similarly, the RNF182-concerted degradation of RELA/p65 reduces toll-like receptor (TLR)-induced innate immune responses^86^ and together, these may offer potential routes for reduced inflammatory phenotypes in the WT cells when compared to *CTSL^−/−^*. The interplay of CTSL and these pro-inflammatory markers may prove useful in discerning the contribution of CTSL to inflammation, and whether deleting or inhibiting CTSL is indeed useful for amelioration of pathogenic factors.

The increased abundance of various ribosomal subunits (RPL22, RPL23A, RPS6, RPS8, RPS1, RPS24) and BOP1 in the WT nuclear-enriched fractions suggests a role for nuclear CTSL in protein translation regulation. This may correlate with increased cell proliferation and cell cycle coordination^87^, as seen with CTSL in breast and colon cancer cells, and neurons^88–90^. Nuclear CTSL is prominent in fibroblasts throughout G1/S phases^37^. Similarly, BOP1 and ribosomal biogenesis are mainly concerted in G1/S phases, with loss of BOP1 arresting cells in the G1 phase^91^. Thus, the presence of nuclear CTSL may facilitate expression of these proteins to promote ribosome biogenesis and cellular proliferation, which may promote cancer progression^92^. We also observed elevated cytoplasmic BOP1 in WT HSC-3 cells (Figure 5C; Table S3), which also links to increased cellular proliferation and migration^93^. As such, the impact of CTSL on cell cycle progression may provide a route to oral cancer cell proliferation.

Beyond broad protein abundance changes, our data also reveal novel putative nuclear substrates for both legumain and CTSL, illustrating their broad impact on the nuclear proteome. The identified nuclear CTSL substrates further support a role for CTSL in cell cycle regulation. One of these was cleavage of MKI67 at R^1738^τA^1739^ which may correlate to its turnover. MKI67 expression is elevated during S/G2/M phases of the cell cycle and undergoes proteasomal degradation in the G1 phase or upon cell cycle exit^94^. It is also upregulated in a range of cancer types, including oral cancer^95^. While we observed numerous dimethylated N-termini for MKI67 in our dataset (Table S4), potentially indicating its degradation, processing at R^1738^τA^1739^ is the only cleavage site enriched in WT cells, suggesting its specific cleavage by CTSL. Whether this event contributes to normal protein turnover of MKI67, or it alters its cellular function is unknown. The most enriched putative cleavage site of CTSL was processing of poly(ADP-ribose) glycohydrolase (PARG) at R^684^τR^685^ (Figure 5F). This site is situated within the catalytic domain (610-795aa) of PARG and would ultimately split the N-terminal NLS (10-16aa) from the C-terminal Poly(ADP-ribose)-binding domain (726-874aa). PARG primarily functions in DNA repair and cell replication^96^. In the context of head and neck squamous cell carcinoma (HNSCC), inhibiting PARG blocks cellular replication and induces cell cycle arrest, resulting in apoptotic pathway stimulation^97^. These cleavage events provide additional support for the role of CTSL in cell cycle progression and regulation, yet the validation and complete functional outcomes of these cleavage events require further investigation.

We also identified the nuclear import protein importin subunit alpha-7 (KPNA6) as a putative CTSL substrate, cleaving at R^13^τM^14^ (Figure 5F). Other importin-α subunits are cleaved following Asp-58 by caspase-3 during apoptosis, and this abolishes their ability to bind chromatin for DNA synthesis^98^. The N-terminal IBB domain of KPNA6 binds to the KPNA6 ARM domain to maintain a closed state which can then open upon binding of the substrate nuclear localisation signal (NLS). KPNA6 can then bind importin-β1 for nuclear translocation. Substrate release in the nucleus and subsequent reformation of the closed state then permits importin recycling back to the cytoplasm^99^. The identified cleavage occurs prior to the basic motif for binding to the ARM domain (^28^RRRR^31^-(X)_17_-^49^KRR^51^). As such, this cleavage may influence autoinhibition of KPNA6 by either promoting a closed state and preventing binding to substrates for nuclear import or may facilitate a constitutively open state to upregulate import. Considering KPNA6 contributes to nuclear import of activated signal transducer and activator of transcription 1 (STAT1) and STAT3, as well as NFκB, further research on the subcellular localisation of these proteins in oral cancer is required.

Overall, our data provide the first systematic analysis of nuclear substrates for the lysosomal proteases CTSL and LGMN. While this research provides insight into the plausible nuclear proteolytic activity of these proteases, it does not confirm the exact location of each cleavage event but rather shows the localisation of the resulting proteolytic products at the time of cell harvest. Mutating either the proteases or their putative substrates to prevent their nuclear translocation will likely be required to discern whether these cleavages occur inside or outside the nucleus. Further, despite identification of numerous neo-N-termini, these require further validation. Employing *in vitro* cleavage assays using recombinant proteins or moving to *in vivo* models of oral cancer would provide more robust results for cross-validation of cleavage events identified in this study. Taken together, this study exemplifies the dependence of legumain in lysosomal cathepsin processing, subsequently influencing the nuclear localisation of CTSL. Our results catalogue a range of protein abundance and proteolytic changes in subcellular compartments of oral cancer cells dependent on CTSL and LGMN and offers novel insights into their proteolytic functions in extra-lysosomal circumstances.

## Methods

### Cell culture

All cells were cultured at 37 °C with 5% CO_2_. RAW264.7 cells (mouse monocyte/macrophage line) were cultured in Dulbecco’s Modified Eagle Medium (DMEM, high glucose, Gibco) supplemented with 10% Foetal Bovine Serum (FBS, CellSera) and 1% antibiotic-antimycotic (100 U/mL penicillin-streptomycin, Thermo). Cells were passaged 1:10 once reaching 80-90% confluence using a cell scraper. HSC-3 cells (human oral squamous cell carcinoma line derived from metastatic lymph node) were grown in DMEM containing 10% FBS and 1% antibiotic-antimycotic. SCC-9 cells (human oral squamous cell carcinoma line derived from tongue) were cultured in DMEM/F12 (Gibco) containing 10% FBS, 1% antibiotic-antimycotic, and 400 µg/mL hydrocortisone (Sigma). Cells were lifted with trypsin (0.25% for HSC-3 and 2.5% for SCC-9) and passaged at 1:10.

### Mice

*Lgmn*^−/−^ C57BL/6N mice were kindly provided by Thomas Reinheckel^100^ and bred in the laboratory of Brian Schmidt and Nigel Bunnett at New York University. Studies were approved by and carried out in accordance with the NYU Institutional Animal Care and Use Committee. Splenic tissues were harvested from 8-week-old healthy male mice (WT and *Lgmn^−/−^*), snap frozen and stored at –80 °C. Spleens were lysed by sonication in citrate buffer (50 mM citrate (Thermo, pH 5.5), 0.5% CHAPS (Sigma), 0.1% Triton X-100, 4 mM DTT (Sigma)), and solids were cleared by centrifugation (21,000 x *g*, 5 minutes, 4 °C).

### Activity-based probe labelling

For live-cell-labelling experiments, cells were plated in 6-well plates to 80% confluence. Once adhered, activity-based probes (ABPs) for legumain^55^ (LE28) or cathepsins^57^ (BMV109) were added (1 µM final, diluted from a 1 mM DMSO stock; 0.1% DMSO final) and incubated for 4h. Cells were washed with PBS, scraped from the wells, and lysed on ice in PBS containing 0.1% Triton X-100. Solids were cleared by centrifugation (21,000 x *g*, 5 minutes, 4 °C) and a bicinchoninic acid (BCA) assay (Pierce) was used to determine total protein concentration. For post-lysis assessment of protease activity, cells were alternatively lysed in citrate buffer (as above) to maintain protease activity. Total protein (80 µg from cell or splenic lysate) was diluted in 20 µL citrate buffer and ABPs were added (1 µM, diluted from a 100 µM DMSO stock; 1% DMSO final) and incubated at 37 °C for 30 min. For both live-cell and lysate labelling, proteins were solubilised by addition of 5x sample buffer (50% glycerol (Sigma), 250 mM Tris-Cl (Sigma), pH 6.8, 10% SDS (VWR LifeSciences), 0.04% bromophenol blue (Sigma), 6.25% beta-mercaptoethanol (Sigma), diluted to 1x final). Samples were boiled at 95 °C for 5 min, and proteins were resolved on a freshly prepared 15% SDS-PAGE gel. To detect ABP binding, gels were scanned with a Cy5 filter set on a Typhoon 5 flatbed laser scanner (GE Healthcare).

### Cell fractionation

Cells were seeded in 10-cm dishes. When confluent, cells were collected and lysed with 200 μL cytoplasmic lysis buffer (10 mM HEPES, 1.5 mM MgCl_2_, 10 mM KCl, 0.5 mM DTT, and 0.05% NP40) followed by centrifugation at 4 °C, 3000 rpm for 10 min. The supernatant containing the cytoplasm-enriched fraction was collected and the pellets washed twice with 200 µL cytoplasmic lysis buffer. The remaining pellets containing nuclear-enriched fractions were solubilised with 50 μL nuclear lysis buffer (5 mM HEPES, 1.5 mM MgCl_2_, 0.2 mM EDTA, 0.5 mM DTT, 26% glycerol (v/v), and 300 mM NaCl), followed by sonicating twice at 30% amplitude (QSONICA Q500) for 5 sec. The homogenate was left on ice for 30 min, followed by centrifugation at 21,000 x *g* for 20 min at 4 °C. Supernatants containing the nuclear-enriched fraction were collected. For cytoplasmic proteins, an equal protein amount (in general 80 µg) was resolved on homemade 15% SDS-PAGE gels. The nuclear-enriched protein samples were loaded at the same volume as their corresponding cytoplasmic-enriched samples and were 4 times more concentrated. The purity of the cytoplasm– and nuclear-enriched fractions was assessed by blotting for tubulin and lamin-A, respectively, as below.

### Immunoblotting

Proteins were transferred from gel to nitrocellulose membranes using the Trans-Blot Turbo Transfer system (Bio-Rad) and incubated with the indicated primary antibody overnight at 4°C. Blots were washed in PBS containing 0.05% Tween-20 (PBST; Sigma) three times before incubation with the secondary antibody for one hour at room temperature and three washes with PBST. A final wash in PBS was performed prior to detection. Horseradish peroxidase (HRP)-conjugated antibodies were detected with Clarity ECL Substrate (Bio-Rad) on a ChemiDoc (Biorad). Fluorophore-conjugated antibodies were visualised using the Typhoon 5 IRlong channel. Ponceau S stain was used to evaluate loading and transfer efficacy. All antibodies were diluted in 1:1 Intercept blocking buffer (LI-COR) and PBST. Antibodies used in this study included goat anti-mouse legumain (1:1,000, R&D AF2058), goat anti-human legumain (1:1,000, R&D AF2199), rabbit anti-β-actin (1:10,000, Life Technologies, MA5-15739), goat anti-mouse cathepsin B (1:1,000, R&D AF965), goat anti-mouse cathepsin L (1:1,000, R&D AF1515), goat anti-mouse cathepsin X (1:1,000, R&D AF1033), goat anti-human cathepsin S (1:500, R&D AF1183), goat anti-mouse cathepsin D (1:1,000, R&D AF1029), goat anti-mouse cathepsin C (1:1,000, R&D AF1034), goat anti-human cathepsin V (1:1,000, R&D AF1080), rabbit anti-alpha tubulin (1:10,000, Abcam ab52866), rabbit anti-lamin A (1:10,000, Invitrogen MA5-35284), donkey anti-goat IgG HRP-conjugated (1:10,000, Novex Life Technologies A15999), donkey anti-goat IgG IR800-conjugated (1:10,000, Li-cor 926-32214), goat anti-rabbit IgG IR800-conjugated (1:10,000, Li-cor 926-32213).

### Immunoprecipitation

Following collection of tissue or cell lysates and BCA as described above, 100 µg protein was aliquoted and labelled with 1 µM BMV109 from a 100x DMSO stock (100 µM) for 30 min at 37°C. Labelling was quenched with addition of 5x sample buffer and the sample split into five equal portions for the input and pulldowns. Immunoprecipitation (IP) buffer (500 µL, PBS, pH 7.4, 0.5% NP-40, 1 mM EDTA) was added to each pulldown sample, along with 10 µL antibody against the protein of interest (goat anti-mouse cathepsin X, goat anti-mouse cathepsin B, goat anti-human cathepsin S, goat anti-mouse cathepsin L) and 40 µL of pre-equilibrated Protein A/G PLUS-Agarose (Santa Cruz, sc-2003) for overnight incubation on a rotating wheel at 4°C. Pulldown samples were washed once in 1 mL IP buffer, followed by 4 times in 0.5 mL IP buffer and a final wash in 0.5 mL 0.9% NaCl before adding 20 µL 2x sample buffer and analysis by SDS-PAGE as described above.

### Secretion analysis and uptake assay

For analysis of secreted proteins, cells were plated in 6-well plates and permitted to adhere. The following day, cells were washed with PBS and media was replaced with 1 mL serum-free media for 24 h. Conditioned media (CM) was collected, centrifuged to pellet cells and debris, and then concentrated using 3 kDa Ultra Centrifugal Filters (Amicon, UFC5003). CM (typically ∼45 µl) was solubilised with 5x sample buffer and resolved on 15% SDS-PAGE gels as above. For the uptake assay, wild-type and legumain-deficient HSC-3 and RAW264.7 cells were seeded in 10 cm dishes with 10 mL complete media. These were left overnight at 37°C with 5% CO_2_ prior to collection of the CM and concentration using 10 kDa Ultra Centrifugal Filters (Amicon, UFC9010). The resulting concentrate was added to legumain-deficient HSC-3 and RAW264.7 cells seeded in a 6-well plate and topped up to 2 mL total volume media. After 48 h, cells were collected for activity-based probe lysate labelling and immunoblotting as described above.

### Legumain and cathepsin L deletion by CRISPR/Cas9

The CRISPR/Cas9 system was used to generate legumain-deficient RAW264.7, HSC-3 and SCC-9 cells, as well as cathepsin L-deficient HSC-3 cells. For RAW264.7 cells, a murine legumain guide RNA (gRNA) construct was used to target exon 8 of the legumain gene (Table 1). As a control, RAW264.7 cells were also transfected with gRNA targeting the human BCL2-like gene, *hBIM*, which is not found in the mouse genome. For human cell lines, HSC-3 and SCC-9, a human legumain gRNA construct was used to target exon 4 of the legumain gene. The oligoduplex was ligated into a SpCas9-(BB)-2A-GRP plasmid (NovoPro V012526) and transfected by nucleofection for RAW264.7 cells, or use of X-tremeGENE 9 DNA Transfection reagent (Roche, XTG9-RO) for HSC-3 and SCC-9 cells. Cells were left to grow in a 6-well plate for 24 hours before single-cell sorting of GFP-positive cells using a BD Influx at the Murdoch Children’s Research Institute (MCRI Melbourne, Australia). The presence of legumain in each single-cell population was determined by activity-based probe lysate labelling with LE28 and western blotting as above, as well as sequencing and Inference of CRISPR Edits (ICE) analysis (Table 2). Further details can be found in the supplementary information.

**Table 1.**
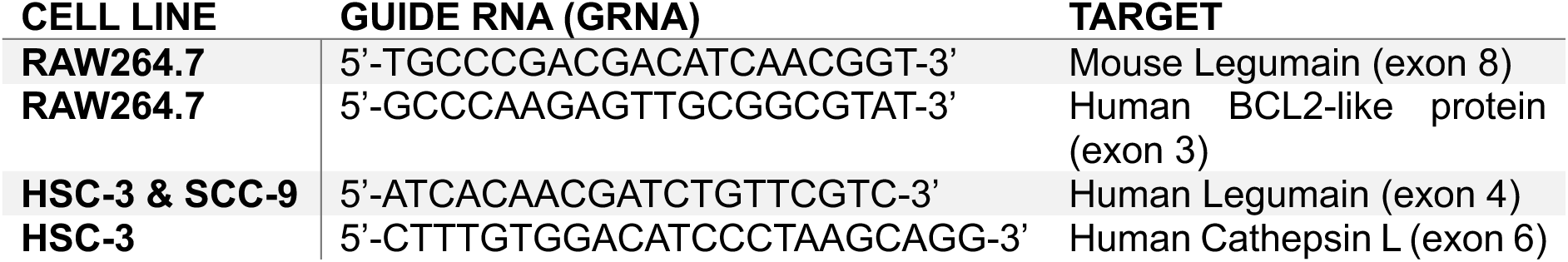
gRNA used for CRISPR/Cas9 knockout of legumain and cathepsin L in various cell types.

**Table 2.**
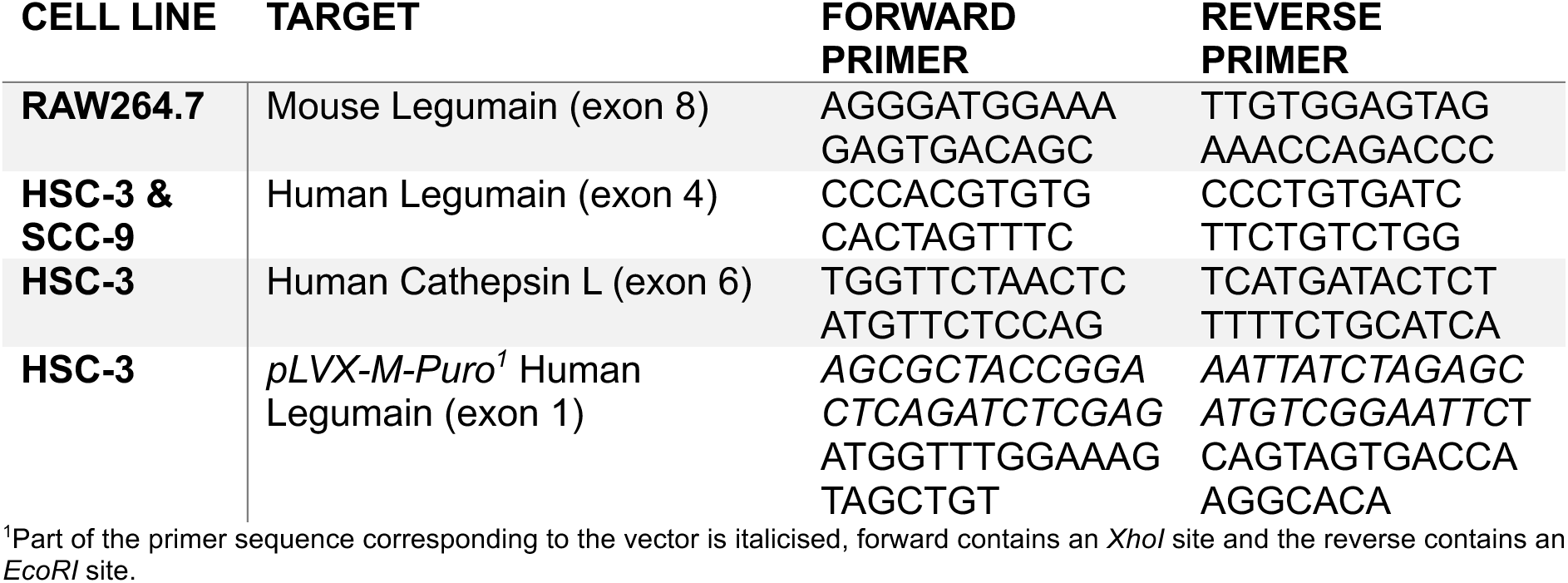
Primers used for re-expression of legumain in HSC-3 and sequencing of CRISPR cut sites for Inference of CRISPR Edits (ICE) analysis.

### Legumain re-expression

The Puc57 vector containing three different constructs of the human legumain cDNA were purchased from Biomatik: WT legumain, catalytically dead mutant legumain (C189S), and integrin-binding mutant legumain (R118H). These constructs were amplified by Q5 high-fidelity PCR using the primers below (Table 2) to generate sequences with overhangs for subsequent *XhoI* and *EcoRI* cleavages and Gibson assembly. The purified PCR products were cloned into the retroviral pLVX-M-Puro vector^101^ (Plasmid #125839) by Gibson assembly as instructed by the manufacturer (NEB E5510). The plasmids containing legumain cDNA, as well as the pGag-pol and pVSV-G vectors were transfected into HEK293T cells using Lipofectamine 3000 reagent (Invitrogen, L3000015) according to manufacturer’s instructions. The supernatant produced was then collected and cleared at 2,000 x *g* for 5 min, prior to filtering through 0.45 µm syringe filters (Millipore Millex-HV PVDF membrane, SLHVR33RB). The resulting viral supernatant was then added to *LGMN^−/−^* HSC-3 cell populations in a dropwise manner along with 8 µg/mL polybrene transfection reagent (Sigma TR-1003) for 24 h. Successfully transduced cells were selected with 1 µL/mL puromycin (Gibco, A1113802) and surviving cells were grown for biochemical analyses.

### In vitro cleavage assays

Recombinant cathepsins were incubated with activated recombinant human legumain (1.5 µg/µL stock, 0.025 µg/µL final concentration) at 1:1 mass ratio (0.5 µg each) in acetate buffer (50 mM sodium acetate (ChemSupply), 100 mM NaCl (EMSURE)) at pH 4.5, 5.5, or 6.5, and incubated at 37 °C for 5 h. Samples in the absence of legumain were used as a negative control. The reactions were quenched by addition of 5x sample buffer (1x final concentration) prior to analysis by SDS-PAGE. Band visualisation was achieved using 0.1% Coomassie brilliant blue G-250 dye (Biorad) in 50% methanol, 10% acetic acid. Briefly, gels were stained in Coomassie solution for 30 min at room temperature with shaking followed by three rounds of destaining in 30% ethanol, 10% methanol for 10 min each. Gels were rinsed in Milli-Q water overnight prior to imaging on the Typhoon 5 IRlong channel. Recombinant proteins used in this study included recombinant human legumain (gifted by Hans Brandstetter)^102^, recombinant mouse cathepsin B (R&D, 965-CY), recombinant human cathepsin L (R&D, 952-CY), recombinant human cathepsin D (R&D, 1014-AS), recombinant human cathepsin X/Z (R&D, 934-CY), and recombinant trypsinogen from bovine pancreas (Sigma, T1143).

### Assessment of the proteome and N-terminome by No-enrichment Identification of Cleavage Events (NICE)

Cytoplasmic– and nuclear-enriched fractions were prepared according to the fractionation protocol above, with the addition of Roche cOmplete, EDTA-free protease inhibitor to the lysis buffers. A total of 100 µg protein was used according to BCA results.

Proteins were reduced with 20 mM DTT (80 °C, 10 min, 500 rpm) and alkylated with 50 mM iodoacetamide (37 °C, 30 min, 500 rpm) in the dark followed by quenching with 50 mM DTT (37 °C, 20 min, 500 rpm). Paramagnetic beads (Sera-Mag SpeedBeads 45152105050250 and 65152105050250, GE Healthcare) were prepared by mixing in a 1:1 ratio and washing three times in Milli-Q water before adjusting to a final concentration of 50 µg/µL in Milli-Q water as previously outlined^103^. Conditioned paramagnetic SP3 beads were added to samples (2 mg of SP3 beads, final protein-to-SP3 bead ratio of 1:20) and protein aggregation was initiated by the addition of ethanol (80% final concentration). Samples were then gently shaken (25 °C, 1,000 rpm) for 20 min prior to washing three times with 500 µL of 80% ethanol using a magnetic rack and resuspending in 90 µL of 200 mM HEPES (pH 7.5). Proteins were dimethylated by adding 30 mM formaldehyde (Sigma) and 30 mM sodium cyanoborohydride (Sigma) and shaking (37 °C, 1,000 rpm) for 1 h. This was repeated once more with an additional 30 mM of formaldehyde and 30 mM sodium cyanoborohydride before labelling was quenched by adding 25 µL 4 M Tris-base (pH 6.8) and shaking (37 °C, 1,000 rpm) for 1 h. Excess formaldehyde and sodium cyanoborohydride were removed from samples using SP3 clean up (1 mg SP3 beads; final protein-to-SP3 bead ratio of 1:10) and proteins were precipitated with ethanol (80% final concentration). Samples were gently shaken (25 °C, 1,000 rpm) for 20 min and then washed three times with 500 µL of 80% ethanol using a magnetic rack. SP3 beads were then resuspended in 100 μL of 200 mM HEPES (pH 7.5) and digested overnight at 37 °C with Solu-trypsin (2 µg solu-trypsin, Sigma, trypsin:protein ratio 1:50). The resulting peptide mixtures were collected using a magnetic rack, acidified with Buffer A* (0.1% trifluoroacetic acid, 2% acetonitrile) and desalted using C_18_ StageTips (Empore™, 3M) with the addition of Oligo-R3 resin reverse phase material (Thermo, 113390) as previously described^104^. Samples were dried using a speedvac and stored at –20 °C until analysis.

### Liquid chromatography and tandem mass spectrometry analysis

Proteome samples were re-suspended in Buffer A* and separated using a Vanquish Neo HPLC coupled to an Orbitrap Astral mass spectrometer (Thermo Fisher Scientific) with an Acclaim^TM^ PepMap^TM^ 100 C18 HPLC trap column (20 mm length x 75 µm diameter, 3 µm particle size) and a µPAC Neo high throughput analytical column (5.5 cm length x 75 µm diameter) (all Thermo Fisher Scientific). For each sample comparing wild-type and legumain-deficient HSC-3 cell fractions, ∼200 ng of peptide mixtures was separated using two 30-minute analytical runs, each undertaken with different Orbitrap MS scan ranges (380-980 m/z or 920-1,500 m/z). Whereas samples comparing wild-type and CTSL-deficient HSC-3 cell fractions (200 ng) were separated using one 30-minute analytical run undertaken with the Orbitrap MS scan ranges set between 380-980 m/z. Each 30-minute run was collected by altering the buffer composition from 3% Buffer B (0.1% formic acid, 77.9% acetonitrile, 2% DMSO) to 6% B over 22.7 min, then from 6% B to 23.5% B over 3.7 min, then from 23.5% B to 40% B over 2 min, and further to 50% for 0.1 min. The column was then washed by holding the composition at 99% B for 0.5 min, and increasing the flow to 2 µL/min for the final 0.7 min. Data-independent acquisition (DIA) was undertaken with a single Orbitrap MS scan (resolution of 120k with the Automated Gain Control (AGC) set to a maximum of 500% or maximum injection time of 5 ms) followed by data-independent acquisition (DIA) of MS2 spectra using the Astral (299 scan events, HCD normalized collision energy of 27%, window overlap of 0 m/z, isolation window of 2 m/z, normalised AGC set to 500%, maximum injection time set to 3 ms).

Raw data files were processed and searched using MSFragger (FragPipe v.22.0)^105^ against the unreviewed human proteome (*Homo sapiens,* UniProt Accession: UP000005640, downloaded November 2024, containing 85,024 protein entries), supplemented with common contaminants, and a reverse decoy database (85,024 decoys: 50%). The in-built DIA_SpecLib_Quant workflow was adapted with the following changes for identification and quantification. Data were allowed for cysteine carbamidomethylation as a fixed modification (+57.0215 Da) as well as variable modifications of lysine dimethylation (+28.0313 Da), methionine oxidation (+15.9949 Da), N-terminal acetylation (+42.0106 Da), N-terminal cyclisation (–17.0265/-18.0106 Da), N-terminal dimethylation (+28.0313 Da), and N-terminal lysine dimethylation (+56.0626 Da). Cleavage specificity was set to “SEMI-N_TERM” and “TrypsinR” (Arg-C), allowing a maximum of 2 missed cleavages. Precursor and fragment mass tolerances of 20 ppm and an isotopic error of 3 Da were also included. Protein and peptide-level false discovery rates (FDR) were determined using Philosopher (v.5.0.0) with default settings (FDR threshold set at 1%). Quantification parameters were left as default and performed with DIA-NN (v.1.8.2 Beta)^106^. The resulting outputs were further processed in Perseus (v.1.6.0.7)^107^, removing decoy matches before a log_2_ transformation was applied. Proteins and peptides identified in a minimum of three of four biological replicates in at least one of the groups were selected and missing values imputed based on a downshifted normal distribution (σ-width = 0.3, σ-downshift = –1.8) for statistical analyses. A student’s two-sample t-test was applied for statistical comparison between groups with a significance threshold set to log_2_(fold change) ±1 and –log_10_(p) = 1.3 (p = 0.05). Volcano plots, density plots, pie charts, and Venn diagrams were created using RStudio (v.4.4.2). Gene ontology and pathways analyses were performed using SRplot^108^. N-terminomics data was also processed using TopFINDer^109^ and pLogo^110^ for generation of sequence logos.

### Supporting Information

- Additional methods on the application of CRISPR/Cas9 for genetic knockout in HSC-3, SCC-9, and RAW264.7 cells, (Figure S1) validation data of *LGMN^−/−^* HSC-3 and SCC-9 cells, (Figure S2, S3, and S4) additional data for legumain processing of cathepsin in different cell lines and tissue, (Figure S5 and S6) supporting data for the *in vitro* cleavage assays between legumain and cathepsins, (Figure S7 and S8) supporting data for nuclear localisation of cathepsins, (Figure S9 and S10) validation data of *CTSL^−/−^* HSC-3 cells, (Figure S11 and S12) supporting data for N-terminomics analysis of wild-type and *CTSL^−/−^* cells, (Figure S13) validation data of nuclear legumain in HSC-3 cells, (Figure S14-15) supporting data for N-terminomics analysis of wild-type and *LGMN^−/−^* cells (PDF)
- (Table S1) Single-cell clone sequencing results of knockouts generated for HSC-3, SCC-9, and RAW264.7 cells (PDF)
- (Table S2) Supporting data of proteomic comparison of nuclear-enriched fractions between wild-type (WT) and cathepsin L knockout HSC-3 cells
- (Table S3) Supporting data of proteomic comparison of cytoplasmic-enriched fractions between wild-type (WT) and cathepsin L knockout HSC-3 cells
- (Table S4) Supporting data of N-terminomic comparison of nuclear-enriched fractions between wild-type (WT) and cathepsin L knockout HSC-3 cells
- (Table S5) Supporting data of N-terminomic comparison of cytoplasmic-enriched fractions between wild-type (WT) and cathepsin L knockout HSC-3 cells
- (Table S6) Supporting data of proteomic comparison of nuclear-enriched fractions between wild-type (WT) and legumain knockout HSC-3 cells
- (Table S7) Supporting data of proteomic comparison of cytoplasmic-enriched fractions between wild-type (WT) and legumain knockout HSC-3 cells
- (Table S8) Supporting data of N-terminomic comparison of nuclear-enriched fractions between wild-type (WT) and legumain knockout HSC-3 cells
- (Table S9) Supporting data of N-terminomic comparison of cytoplasmic-enriched fractions between wild-type (WT) and legumain knockout HSC-3 cells

## Acknowledgements

We thank T. Reinheckel for providing access to the legumain-deficient mouse strain and B. Schmidt and N. Bunnett for breeding the mice and providing spleen tissue. We thank E. Dall and H. Brandstetter for the kind gift of recombinant legumain. We thank the Melbourne Mass Spectrometry and Proteomics Facility, the Australian Genome Research Facility at the Peter MacCallum Cancer Centre, and the flow cytometry platform at the Murdoch Children’s Research Institute for assistance in generating data for NICE analysis, single-cell clone sequencing, and single-cell sorting, respectively.

## Author Contributions

**A.R.Z.** Data curation; Formal analysis; Investigation; Methodology; Validation; Visualization; Writing – original draft; Writing – review & editing.

**B.X.** Conceptualization; Data curation; Formal analysis; Investigation; Methodology; Validation; Writing – review & editing.

**B.M.A.** Data curation; Formal analysis; Investigation; Methodology; Project administration; Writing – review & editing.

**L.L.** Data curation; Investigation.

**S.L.** Data curation; Investigation.

**J.S.** Investigation; Methodology.

**D.A.S.** Resources; Supervision.

**N.E.S.** Methodology; Resources; Supervision.

**L.E.E.-M.** Conceptualization; Funding acquisition; Investigation; Methodology; Project administration; Resources; Supervision; Writing – review & editing.

## Data Availability

The mass spectrometry proteomics and N-terminomics data has been deposited in the ProteomeXchange Consortium *via* the PRIDE partner repository with the data set identifiers PDX066582 and PDX067035.

## Abbreviations

ABP: activity-based probe
AEP: asparaginyl endopeptidase
BCA: bicinchoninic acid
CM: conditioned media
CTS: cathepsin
CTSB: cathepsin B
CTSC: cathepsin C
CTSD: cathepsin D
CTSH: cathepsin H
CTSL: cathepsin L
*CTSL^−/−^*: cathepsin L knockout
CTSS: cathepsin S
CTSV: cathepsin V
CTSX: cathepsin X/Z
CUX1: CCAAT-displacement protein/cut homeobox transcription factor 1
DC: double-chain
DIA: data-independent acquisition
ECM: extracellular matrix
EGFR: epithelial growth factor receptor
EMT: epithelial-to-mesenchymal transition
FAIMS: high-field asymmetric waveform ion mobility spectrometry
GO: gene ontology
GRN: progranulin
HSNCC: head and neck squamous cell carcinoma
ICE: inference of CRISPR edits
IL: interleukin
LGMN: legumain
*LGMN^−/−^*: legumain knockout
LMP: lysosomal membrane permeability
M6P: mannose-6-phosphate
MMP2: matrix metalloprotease 2
NICE: no-enrichment identification of cleavage events
NLS: nuclear localisation signal
PTM: post-translational modification
SC: single-chain
STAT: signal transducer and activator of transcription
TGFB: transforming growth factor beta
TLR: toll-like receptor
TME: tumour microenvironment
TNF-α: tumour necrosis factor alpha
TPG: trypsinogen
WT: wild-type

## For Table of Contents Only

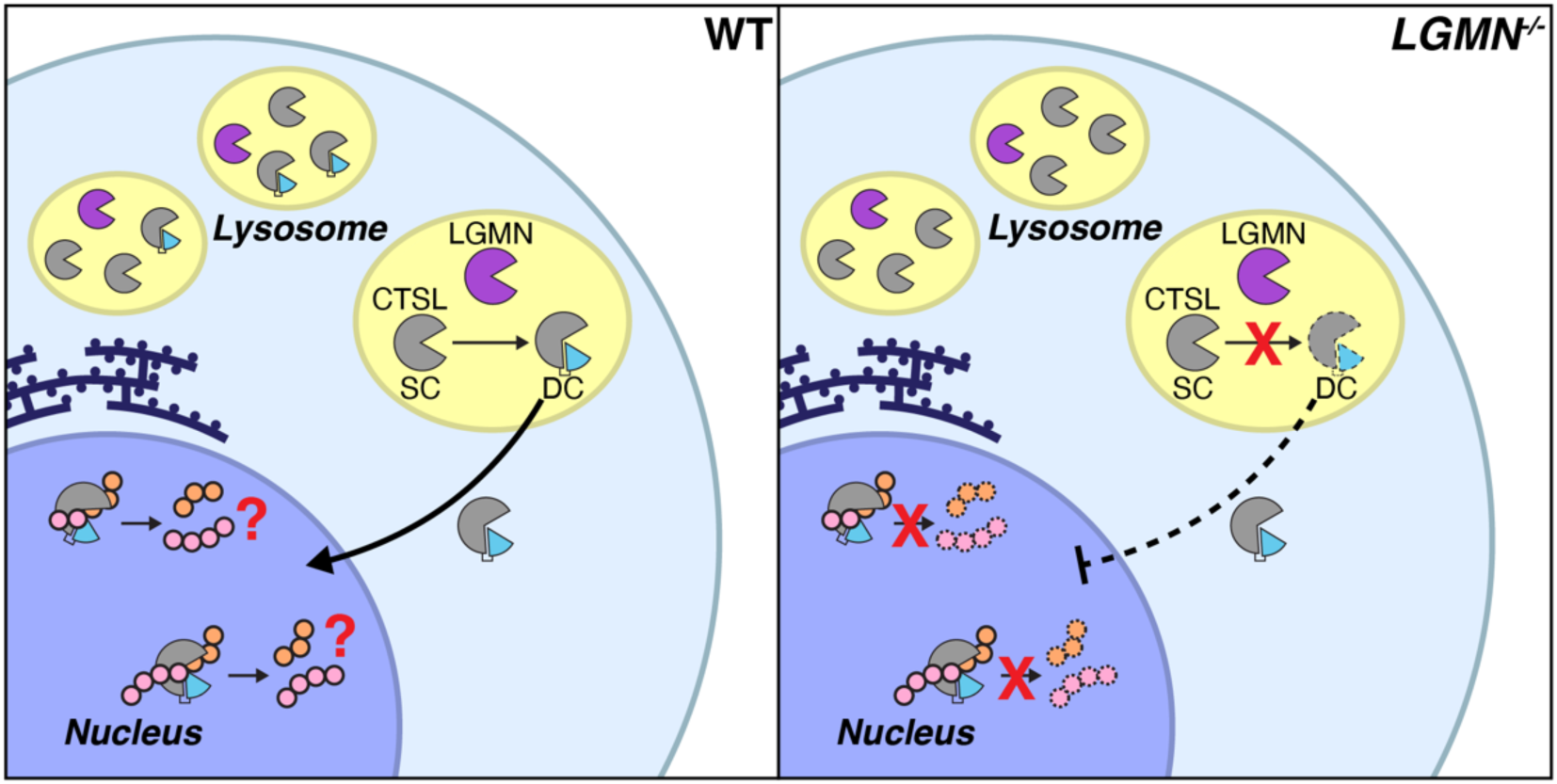

