## Supplemental Figures for "Legumain drives processing of cathepsins and nuclear localisation of cathepsin L"

**Supplemental Information**

### Supplemental Methods

#### *Legumain and cathepsin L deletion by CRISPR/Cas9*

The CRISPR/Cas9 system was used to generate legumain-deficient RAW264.7, HSC-3 and SCC-9 cells, as well as cathepsin L-deficient HSC-3 cells. For RAW264.7 cells, a murine legumain guide RNA (gRNA) construct was used to target exon 8 of the legumain gene (Table 1). As a control, RAW264.7 cells were also transfected with gRNA targeting the human BCL2-like gene, *hBIM*, which is not found in the mouse genome. For human cell lines, HSC-3 and SCC-9, a human legumain gRNA construct was used to target exon 4 of the legumain gene (Table 1). An empty vector was used as control. The gRNA (10 µM final) was ligated to form an oligoduplex by incubating with 10x T4 ligase buffer (1x final, NEB B0202) and T4 polynucleotide kinase (NEB M0201) at 37 °C for 30 min, followed by 95 °C for 5 min and a cycle between 94 °C for 5 sec and 25 °C for 20 sec 70 times. The SpCas9-(BB)-2A-GFP plasmid (NovoPro V012526) was digested with 2 µL *BsbI* (NEB R0539) in 5 µL rCutSmart buffer (NEB B6004) for 1 hour at 37 °C and gel purified (0.8% agarose) using the Monarch DNA gel extraction kit (NEB T1020G). The resulting oligoduplex and digested plasmid were ligated using 10x T4 ligase buffer (1x final) and T4 DNA ligase (NEB M0202) overnight at room temperature.

Competent E. coli cells were transformed with the plasmid complex by heat shock by mixing the for 30 minutes on ice, followed by a 45-second heat shock at 42 °C. E. coli cells were recovered in SOC media (2% (w/v) tryptone, 0.5% (w/v) yeast extract, 8.56 mM sodium chloride, 10 mM magnesium chloride, 10 mM magnesium sulphate, 2.5 mM, potassium chloride and 20 mM glucose [ThermoFisher]) for one hour at 37°C with constant shaking. The E. coli was spread on separate LB-Agar plates containing 100 µg/mL ampicillin and incubated overnight at 37°C. The plasmid was extracted from transformed bacteria using a PureYield Plasmid Miniprep Kit (Promega) and was sequenced at the Australian Genome Research Facility (AGRF, Peter MacCallum Cancer Centre, Melbourne, Australia). Successfully transformed colonies were further grown in LB (1% (w/v) Peptone 140, 0.5% (w/v) yeast extract and 0.5% (w/v) sodium chloride [ThermoFisher]) to amplify the plasmid DNA. DNA was extracted using a PureYield Plasmid Miniprep (Promega) for further transfection.

Transfection of RAW264.7 cells was achieved by Nucleofection. Briefly, cells were aliquoted to contain $2x10^6$ cells per transfection and spun for 10 minutes at 90 *x g*. The supernatant was removed, and the pellet was resuspended in 18 µL nucleofector solution (Nucleofector Kit V, Lonza Bioscience, VCA-1003) with the addition of 2 µL plasmid containing gRNA for either *hBIM* or *mLgmn*. The total volume was transferred to the nucleofector strip with the program for RAW264.7 cells applied. Once complete, pre-warmed complete cell media was added to each transfection reaction and cells were transferred to a 6-well plate for further growth. GFP-positive cells were sorted at Murdoch Children's Research Institute (MCRI Melbourne, Australia) using a BD Influx to obtain single-cell populations. The presence of legumain in each single cell population was checked by activity-based probe lysate labelling with LE28 and western blotting with goat anti-mouse legumain antibody as described above.

Transfection of HSC-3 and SCC-9 cells was performed using the X-tremeGENE 9 DNA Transfection reagent (Roche XTG9-RO). Cells were plated in a 6-well plate and left overnight to reach 70% confluency. Transfection reagent was prepared, containing 100 µL Opti-MEM Reduced Serum Medium (Gibco 31985062), 2 µg of gRNA, and 6 µL X-tremeGENE 9 DNA Transfection Reagent. The solution was left at room temperature for 10 minutes before being added to cells in a dropwise manner. Cells were left to incubate at 37°C with 5% $CO_2$ for 24 hours to allow transfection, before single-cell sorting as above.

Sequencing of single cell clone populations for Inference of CRISPR Edits (ICE) analysis was performed to confirm genetic knockout of legumain or cathepsin L in each respective cell type. Briefly, 200,000 cells were washed in PBS and pelleted for extraction of genomic DNA using 50 µL QuickExtract solution (Biosearch Technologies, QE09050). This was achieved by incubating samples at 65°C for 15 minutes, followed by 68°C for 15 minutes, and then 98°C for a further 10 minutes in a thermocycler. Extracted genomic DNA was amplified using primers flanking the Cas9 cut site (Table 2) and purified using the Wizard Gel Purification Kit (Promega, A9281). The concentration of the purified product was measured by nanodrop and normalised prior to sequencing at the Australian Genome Research Facility (AGRF Melbourne, Australia). The resulting data was analysed using ICE (https://ice.synthego.com/).

### Supplemental Figures

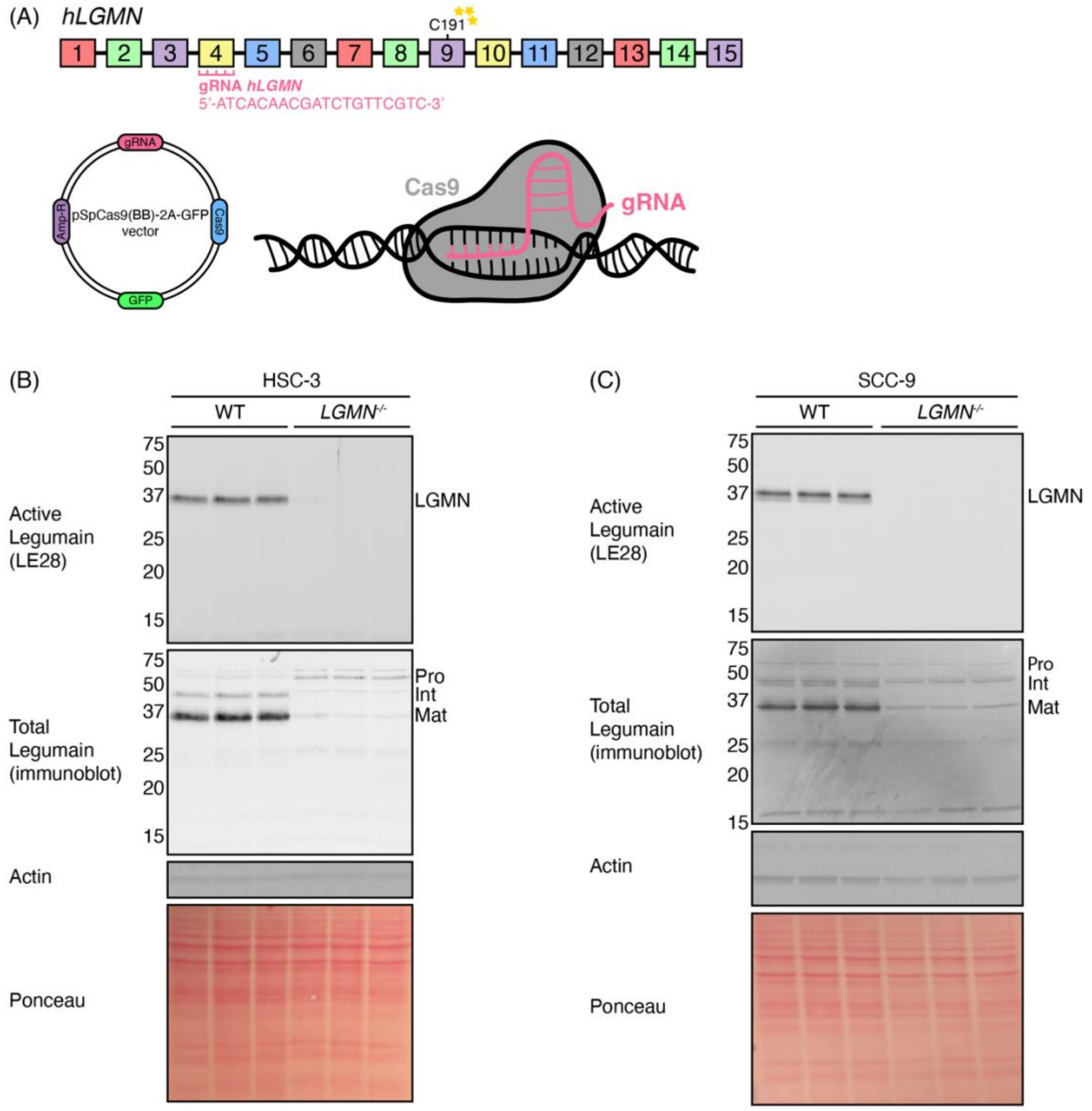


**Figure S1. CRISPR/Cas9 knockout of legumain in human oral cancer cell lines HSC-3 and SCC-9.** (A) Schematic of human legumain (*hLGMN*) knockout by CRISPR/Cas9. (B-C) Lysate labelling of legumain activity using LE28 and subsequent immunblot analysis of wild-type (WT) and legumain-deficient (*LGMN*$^{-/-}$) HSC-3 (B) and SCC-9 cells (C) (n = 3 per group). Actin and Ponceau S stain were used as loading controls.

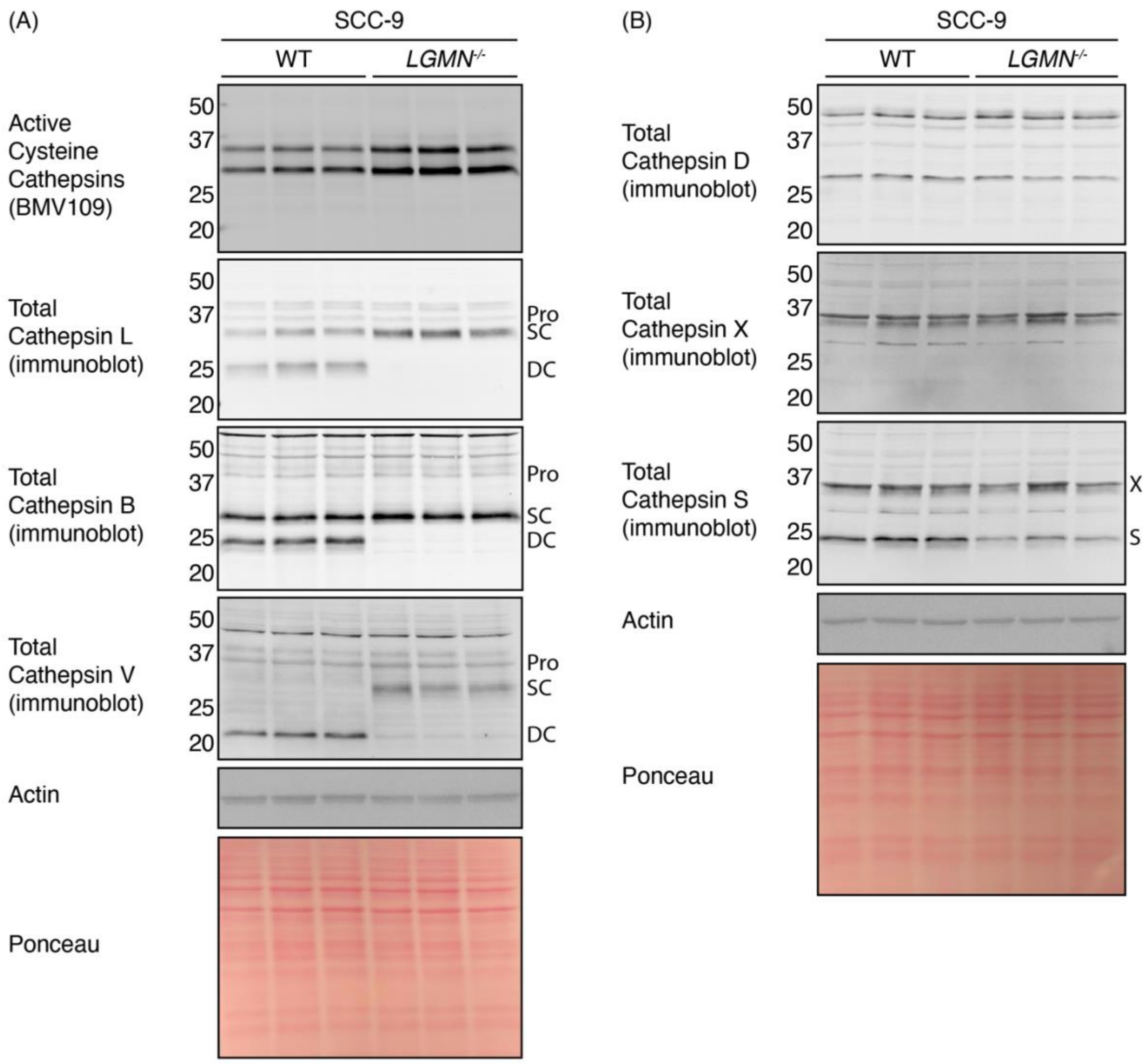


**Figure S2. Lysosomal cathepsins are upregulated in the absence of legumain in SCC-9 cells.** (A) Lysate labelling of wild-type and legumain-deficient (*LGMN*$^{-/-}$) SCC-9 cells with BMV109 to assess cysteine cathepsin activity, and immunoblot of cathepsins L, B, and V (n = 3 per group). (B) Immunoblot analysis of cathepsins D, X, and S (n = 3 per group). Actin and Ponceau S stain are used as loading controls. Cathepsin S immunoblot was performed on the same membrane following cathepsin X immunoblot.

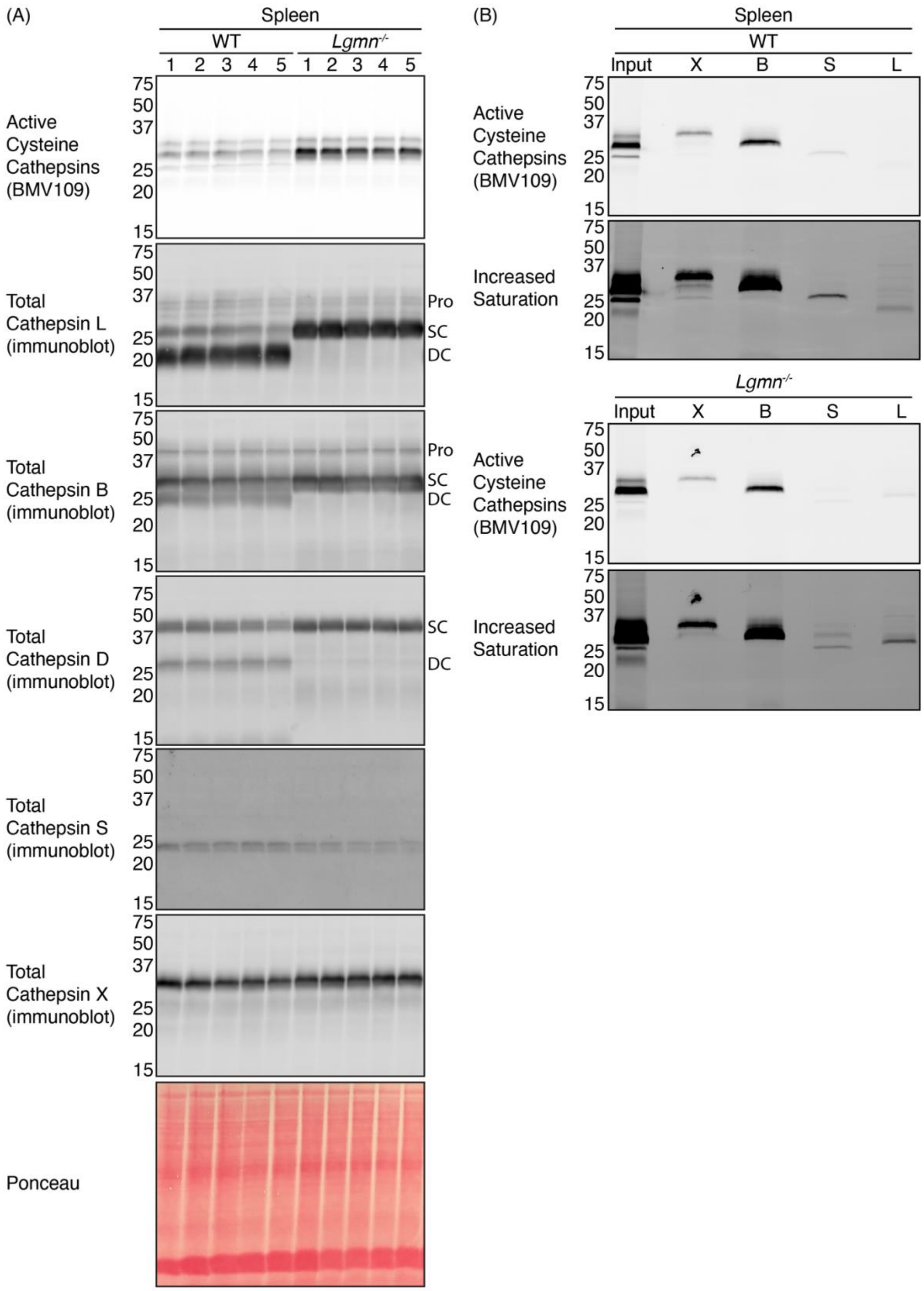


**Figure S3. Lysosomal cathepsins are upregulated in the absence of legumain in murine spleen tissue.** (A) Lysate labelling of wild-type and legumain-deficient (*LGMN*$^{-/-}$) mouse spleen lysates with BMV109 to assess cysteine cathepsin activity, and immunoblot of cathepsins L, B, D, S, and X (n = 5 per group). Ponceau S stain was used as a loading control. (B) Immunoprecipitation of WT and *LGMN*$^{-/-}$ spleen lysates lysate-labelled with BMV109 (n = 3 independent experiments).

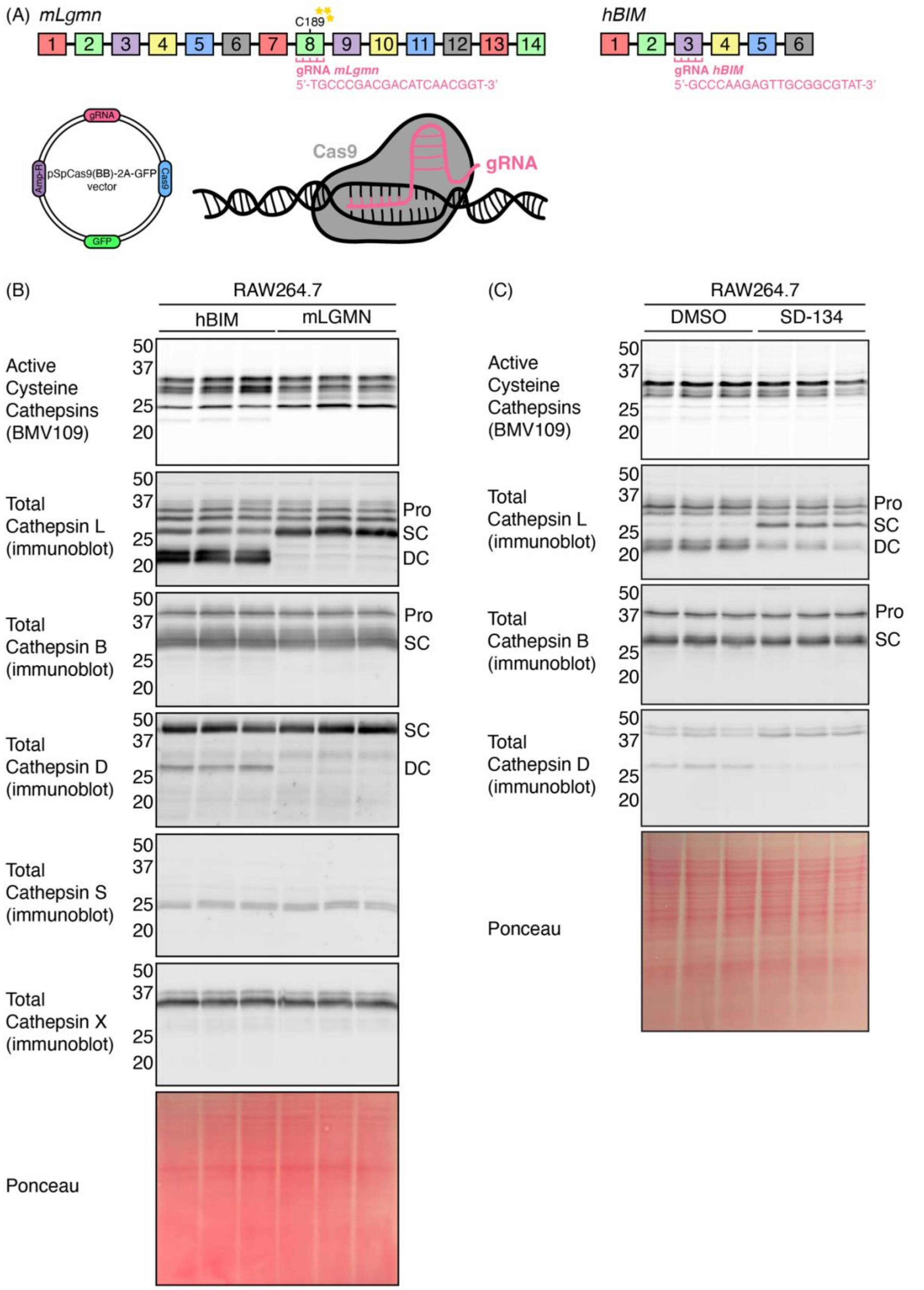


**Figure S4. Legumain depletion in murine macrophages leads to lysosomal cathepsin upregulation dependent on legumain activity.** (A) Schematic of mouse legumain (*mLgmn*) knockout by CRISPR/Cas9 in RAW264.7 cells. (B) Lysate labelling of cysteine cathepsins by

BMV109 in wild-type (hBIM) and legumain knockout (mLGMN) RAW264.7 cells and subsequent immunoblot analysis of cathepsins L, B, D, S, and X (n = 3 per group). (C) Lysate labelling of cysteine cathepsins by BMV109 in RAW264.7 cells treated with 10 μM of a legumain-specific inhibitor (SD-134) or vehicle (DMSO) and subsequent immunoblot analysis of cathepsins L, B, and D (n = 3 per group). Ponceau S stain was used as a loading control.

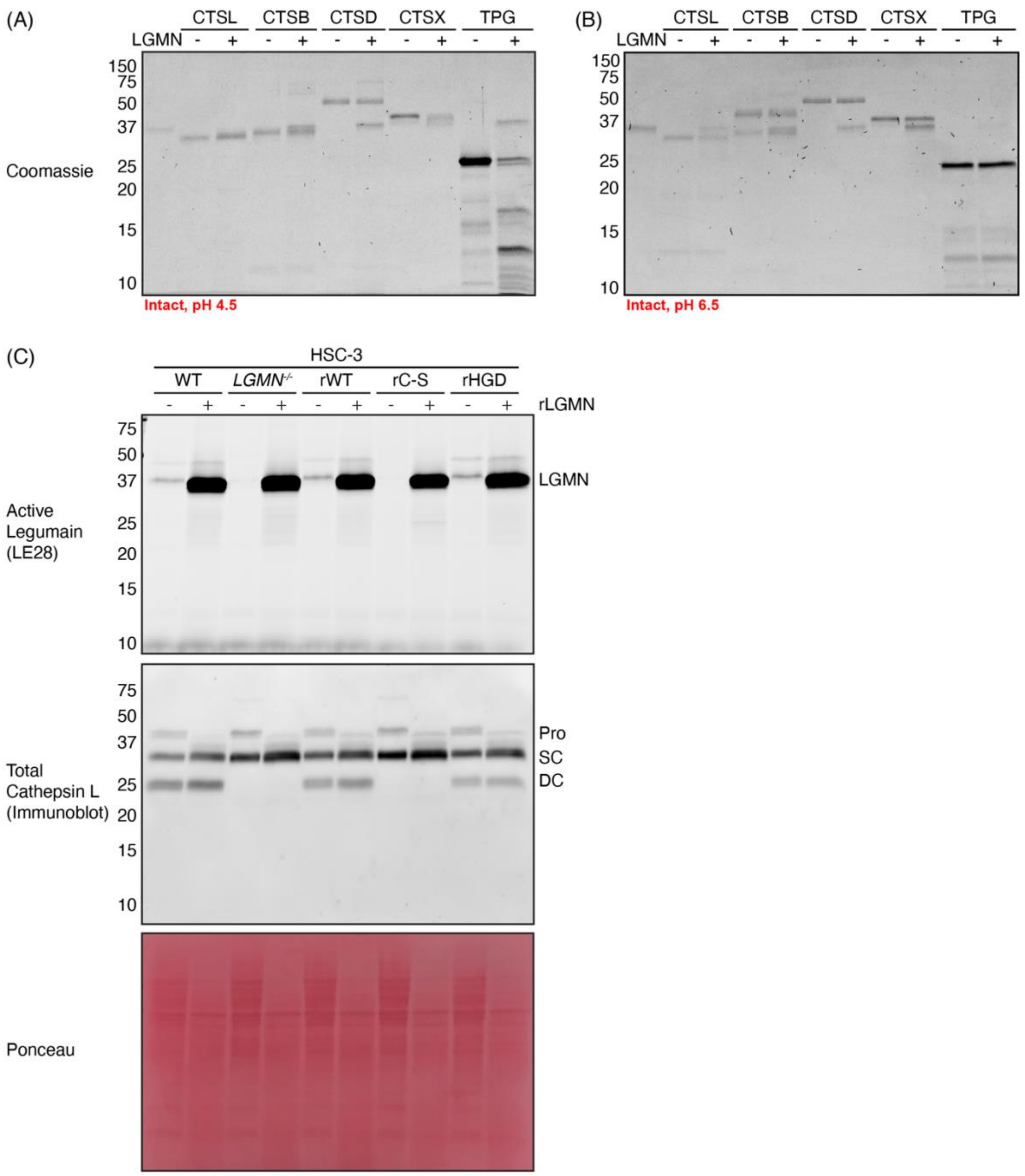


**Figure S5. Addition of recombinant legumain does not induce processing of lysosomal cathepsins *in vitro*.** (A-B) *In vitro* cleavage assay of intact proteins using activated recombinant legumain (LGMN) and recombinant cathepsins L (CTSL), B (CTSB), D (CTSD), and X (CTSX) at pH 4.5 (A) or pH 6.5 (B). Trypsinogen (TPG) was used as a positive control. (C) Treatment of wild-type (WT), legumain-deficient (*LGMN*$^{-/-}$), reconstituted wild-type (rWT), reconstituted catalytically-dead legumain (rC189S, rC-S), and reconstituted integrin-binding mutant legumain (rR118H, rHGD) HSC-3 cell lysates (pH 5.5) with recombinant legumain (rLGMN) and assessment of cathepsin L processing by immunoblot. Ponceau S stain was used as a loading control.

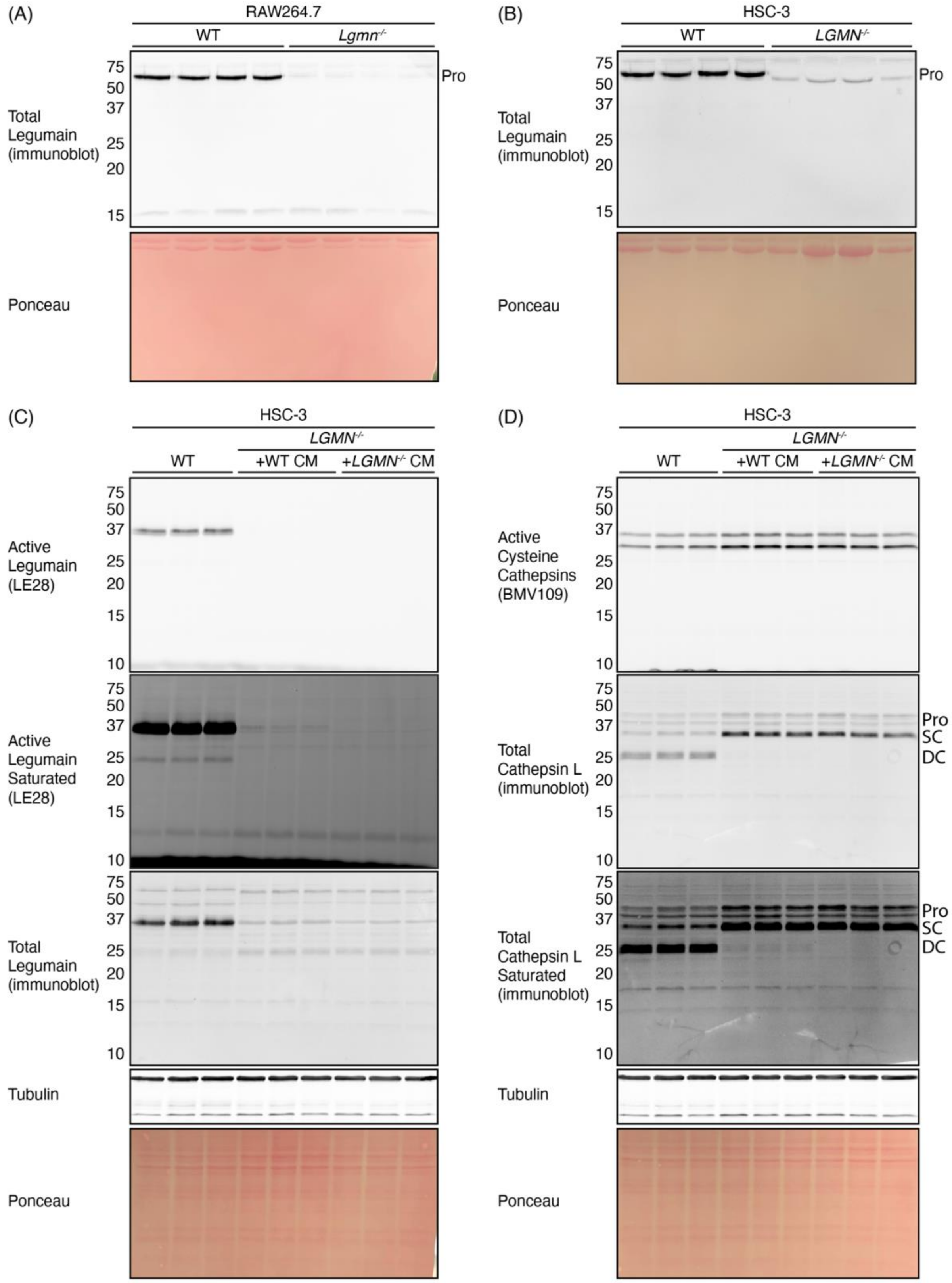


**Figure S6. Uptake of exogenous legumain rescues processing of cathepsin L in legumain-deficient cells.** (A-B) Immunoblot analysis of legumain in conditioned media collected from wild-type (WT) and legumain-deficient (*LGMN*$^{-/-}$) RAW264.7 (A) and HSC-3 cells (B). Ponceau S stain was used as a loading control (n = 4 per group). (C) Assessment of legumain uptake and activity by lysate labelling with LE28 and immunoblot analysis in wild-type (WT) cells and legumain-deficient (*LGMN*$^{-/-}$) HSC-3 cells treated with conditioned media (CM) from WT or *LGMN*$^{-/-}$ cells (n = 3 per group). (D) Lysate labelling of cysteine cathepsins with BMV109 and analysis of cathepsin L expression by immunoblot analysis in wild-type (WT) cells and legumain-deficient (*LGMN*$^{-/-}$) HSC-3 cells treated with conditioned media (CM) from WT or *LGMN*$^{-/-}$ cells (n = 3 per group). Tubulin and Ponceau S stain were used as loading control.

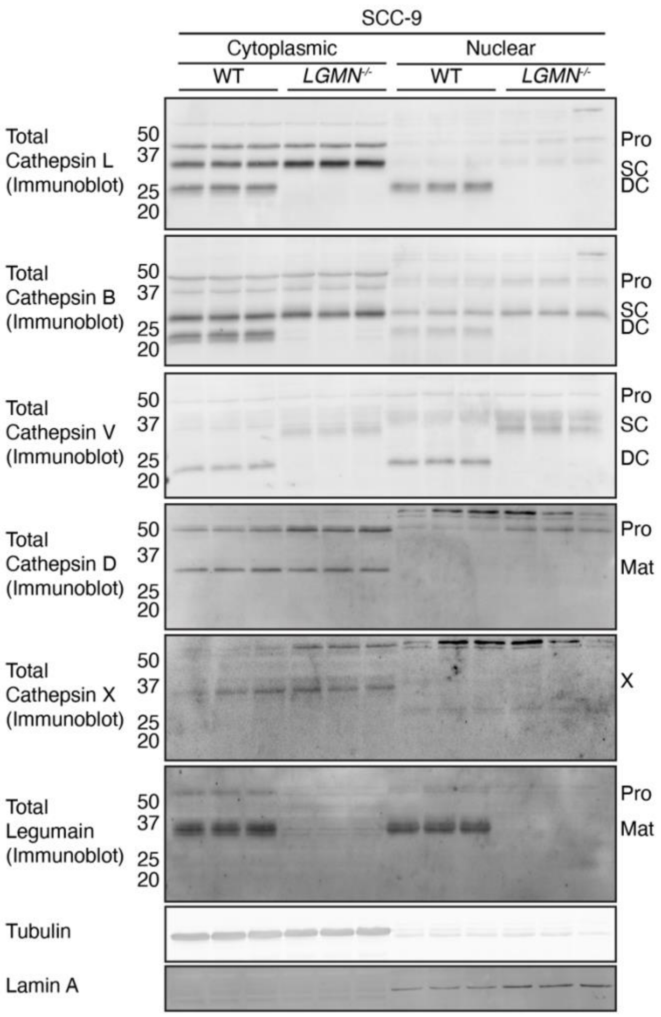


**Figure S7. Cathepsin L nuclear localisation is altered upon loss of legumain in SCC-9 oral cancer cells.** Immunoblot analysis of cathepsins L, B, V, D, and X, and legumain in SCC-9 fractions following subcellular fractionation into cytoplasmic and nuclear-enriched lysates (n = 3 per group). Tubulin and lamin-A were used as markers for cytoplasmic and nuclear fractions, respectively.

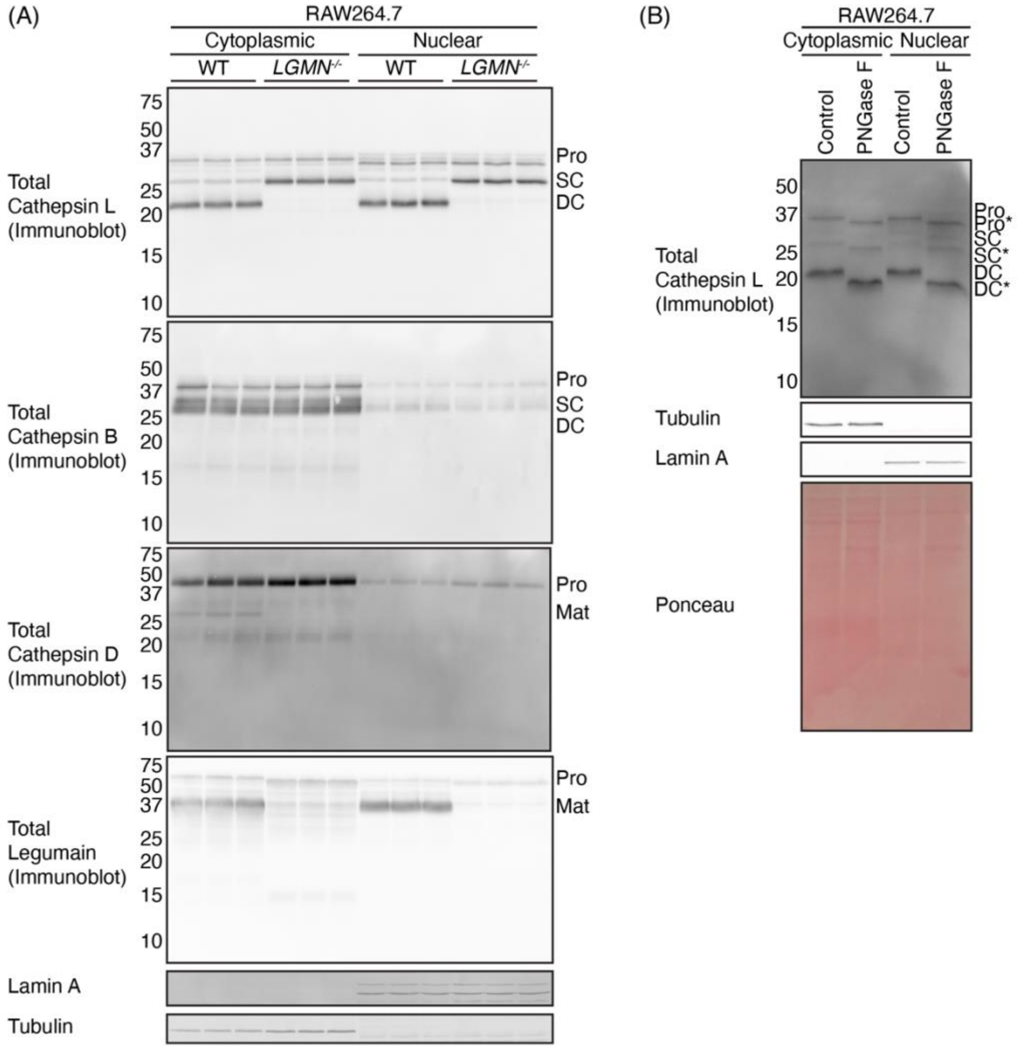


**Figure S8. Changes in the nuclear localisation of lysosomal cathepsins is negligible upon loss of legumain in murine macrophages.** (A) Immunoblot analysis of cathepsins L, B, and D, and legumain in RAW264.7 fractions following subcellular fractionation into cytoplasmic and nuclear-enriched lysates (n = 3 per group). (B) Immunoblot analysis of cathepsin L following treatment of RAW264.7 cytoplasmic-enriched and nuclear-enriched fractions with PNGase F to assess glycosylation. Tubulin and lamin-A were used as markers for cytoplasmic and nuclear fractions, respectively. Ponceau S stain was used as a loading control.

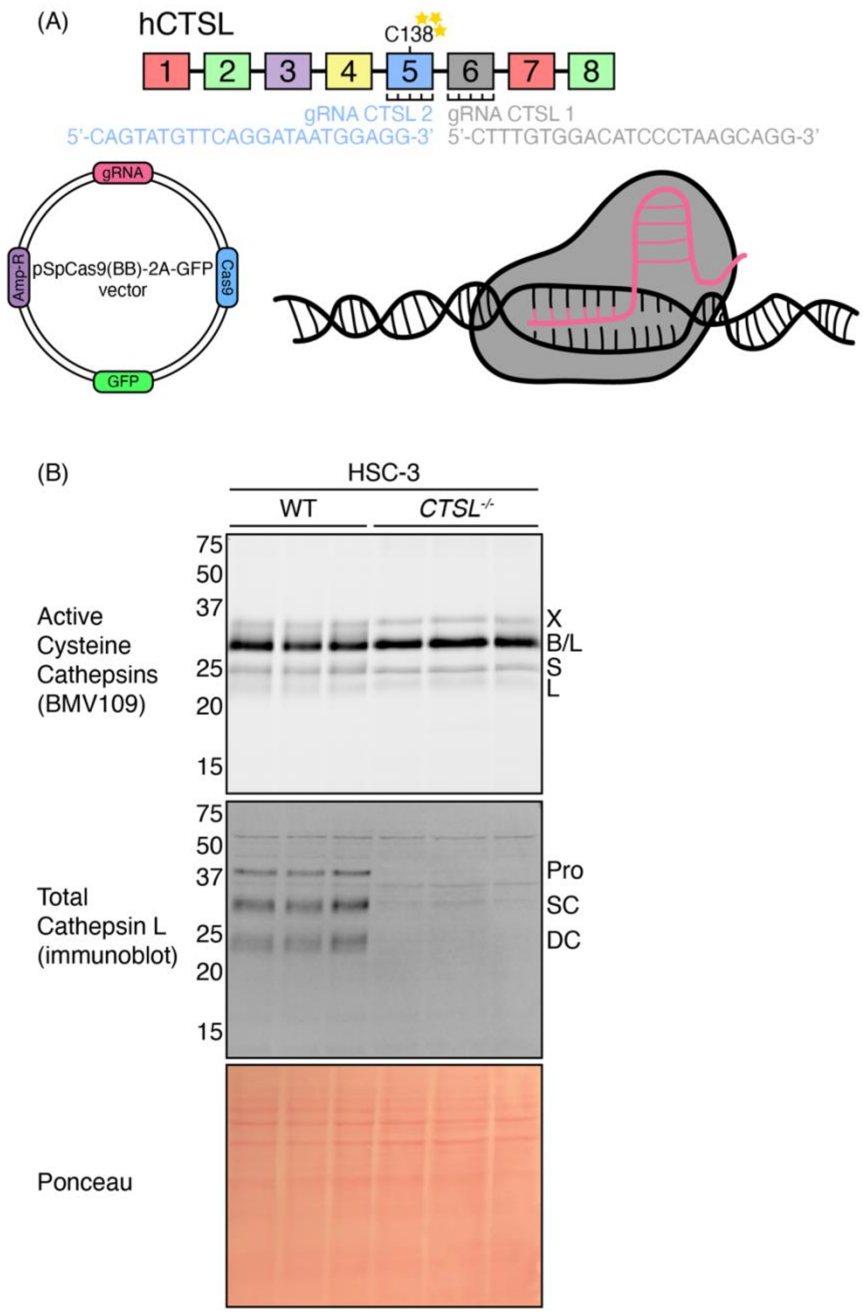


**Figure S9. Cathepsin L knockout in HSC-3 using the CRISPR/Cas9 system.** (A) Schematic of CRISPR/Cas9 depletion of human cathepsin L (*hCTSL*) in HSC-3 cells. (B) Validation of cathepsin L loss by live labelling of wild-type (WT) and cathepsin L-deficient (*CTSL*$^{-/-}$) HSC-3 cells and subsequent immunoblot analysis (n = 3 per group). Ponceau S stain was used as a loading control.

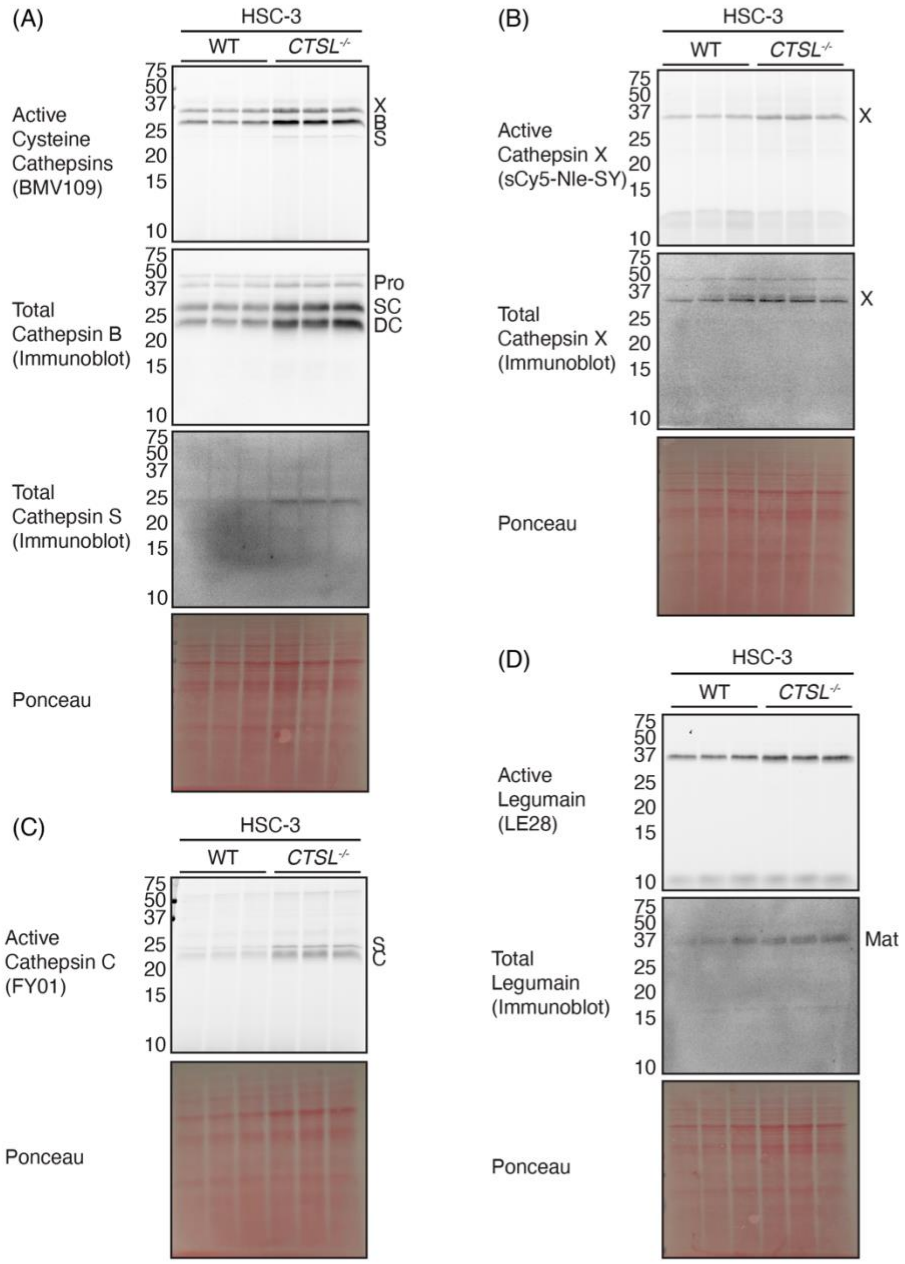


**Figure S10. Lysosomal cathepsins are elevated in the absence of cathepsin L in HSC-3 oral cancer cells.** (A) Lysate labelling of cysteine cathepsins using BMV109 in wild-type (WT) and cathepsin L-deficient (*CTSL*$^{-/-}$) HSC-3 cells, and subsequent immunoblot analysis of cathepsins B and S (n = 3 per group). (B) Lysate labelling of cathepsin X using sCy5-Nle-SY and subsequent immunoblot analysis in WT and *CTSL*$^{-/-}$ cells (n = 3 per group). (C) Lysate labelling of active cathepsin C using FY01 in WT and *CTSL*$^{-/-}$ cells (n = 3 per group). (D) Lysate labelling of active legumain using LE28 and immunoblot analysis for expression in WT and *CTSL*$^{-/-}$ cells (n = 3 per group). Ponceau S stain was used as a loading control.

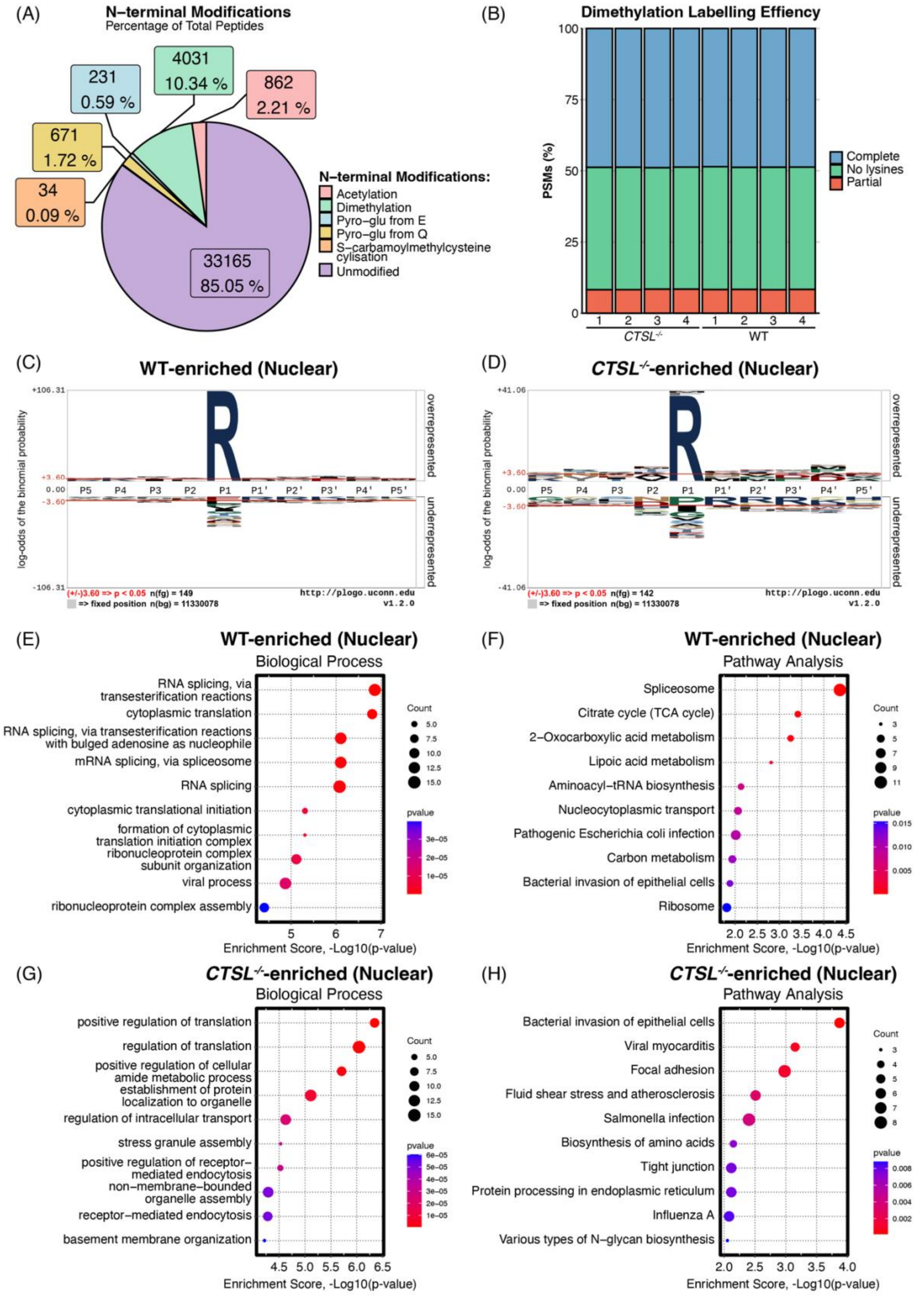


**Figure S11. Bioinformatic analysis of wild-type (WT) and cathepsin L-deficient (*CTSL*$^{-/-}$)-enriched N-termini reveals roles for nuclear cathepsin L in splicing and protein translation regulation.** (A) Proportion of N-terminal modifications of the total peptide identified by mass spectrometry analysis in the No-enrichment Identification of Cleavage Events (NICE) workflow across all biological replicates (n = 4 per group). (B) Dimethylation labelling efficiency in nuclear-enriched fractions collected from WT and *CTSL*$^{-/-}$ HSC-3 cells (n = 4 per group). Peptides with all internal lysines dimethylated represent complete dimethylation (blue), whilst those with lysines missing dimethyl labelling are incomplete (red). Peptides with no internal lysines are also shown (green). (C-D) Sequence logo for the identified cleavage sites in the WT-enriched (C) and *CTSL*$^{-/-}$-enriched (D) N-termini. Figures were created using pLogo with significance indicated by the red line ($p < 0.05$). (E-H) Gene ontology and pathway analysis of WT-enriched (E-F) and *CTSL*$^{-/-}$-enriched (G-H) N-termini using SRplot.

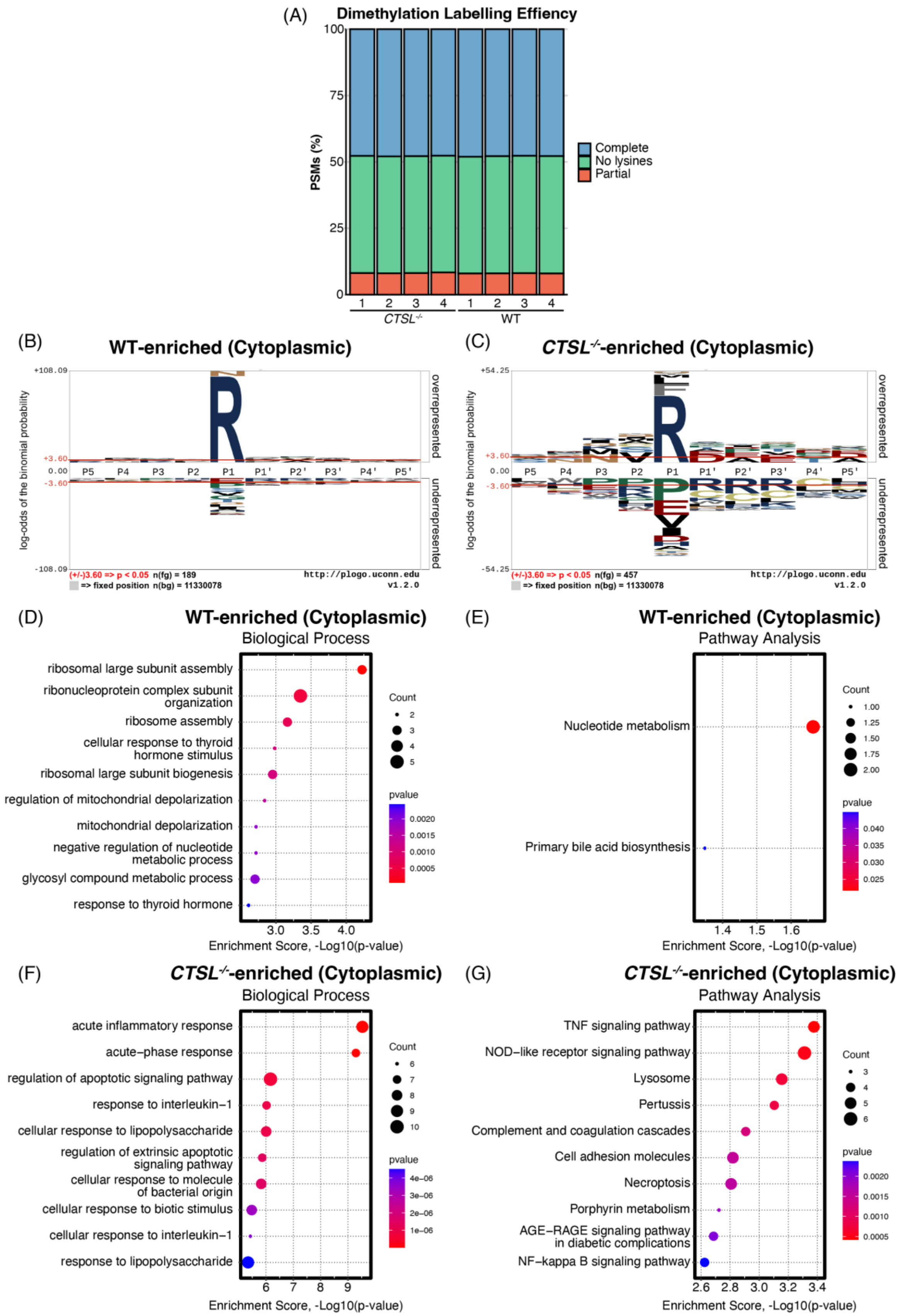


**Figure S12. Bioinformatic analysis of wild-type (WT) and cathepsin L-deficient (*CTSL*$^{-/-}$)-enriched N-termini from cytoplasmic-enriched fractions suggests roles in ribosomal biogenesis.** (A) Dimethylation labelling efficiency in cytoplasmic-enriched fractions collected from WT and *CTSL*$^{-/-}$ HSC-3 cells (n = 4 per group). Peptides with all internal lysines dimethylated represent complete dimethylation (blue), whilst those with lysines missing dimethyl labelling are incomplete (red). Peptides with no internal lysines are also shown (green). (B-C) Sequence logo for the identified cleavage sites in the WT-enriched (B) and *CTSL*$^{-/-}$-enriched (C) N-termini. Figures were created using pLogo with significance indicated by the red line ($p < 0.05$). (D-G) Gene ontology and pathway analysis of WT-enriched (D-E) and *CTSL*$^{-/-}$-enriched (F-G) N-termini using SRplot.

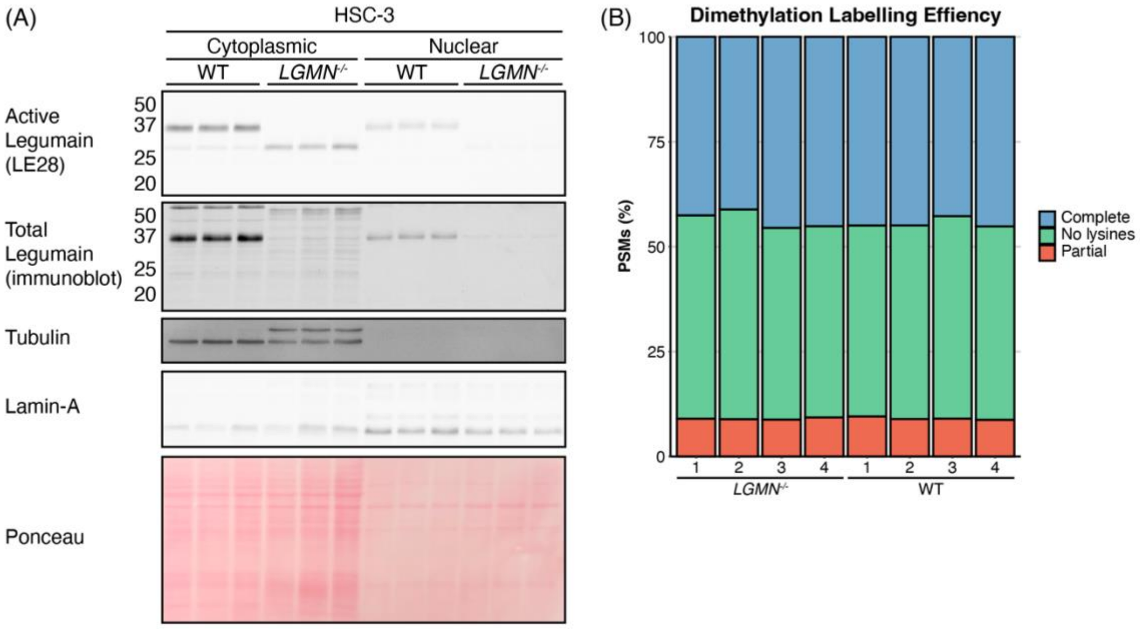


**Figure S13. Legumain is active in nuclear-enriched fractions taken from HSC-3 cells.** (A) Live-labelled HSC-3 cells for analysis of legumain activity by LE28 in cytoplasmic-enriched and nuclear-enriched fractions and immunoblot analysis for legumain expression analysis. Tubulin and lamin-A were used as cytoplasmic and nuclear markers, respectively. Ponceau S stain was used as a loading control. (B) Dimethylation labelling efficiency in nuclear-enriched fractions collected from WT and *LGMN*$^{-/-}$ HSC-3 cells (n = 4 per group). Peptides with all internal lysines dimethylated represent complete dimethylation (blue), whilst those with lysines missing dimethyl labelling are incomplete (red). Peptides with no internal lysines are also shown (green).

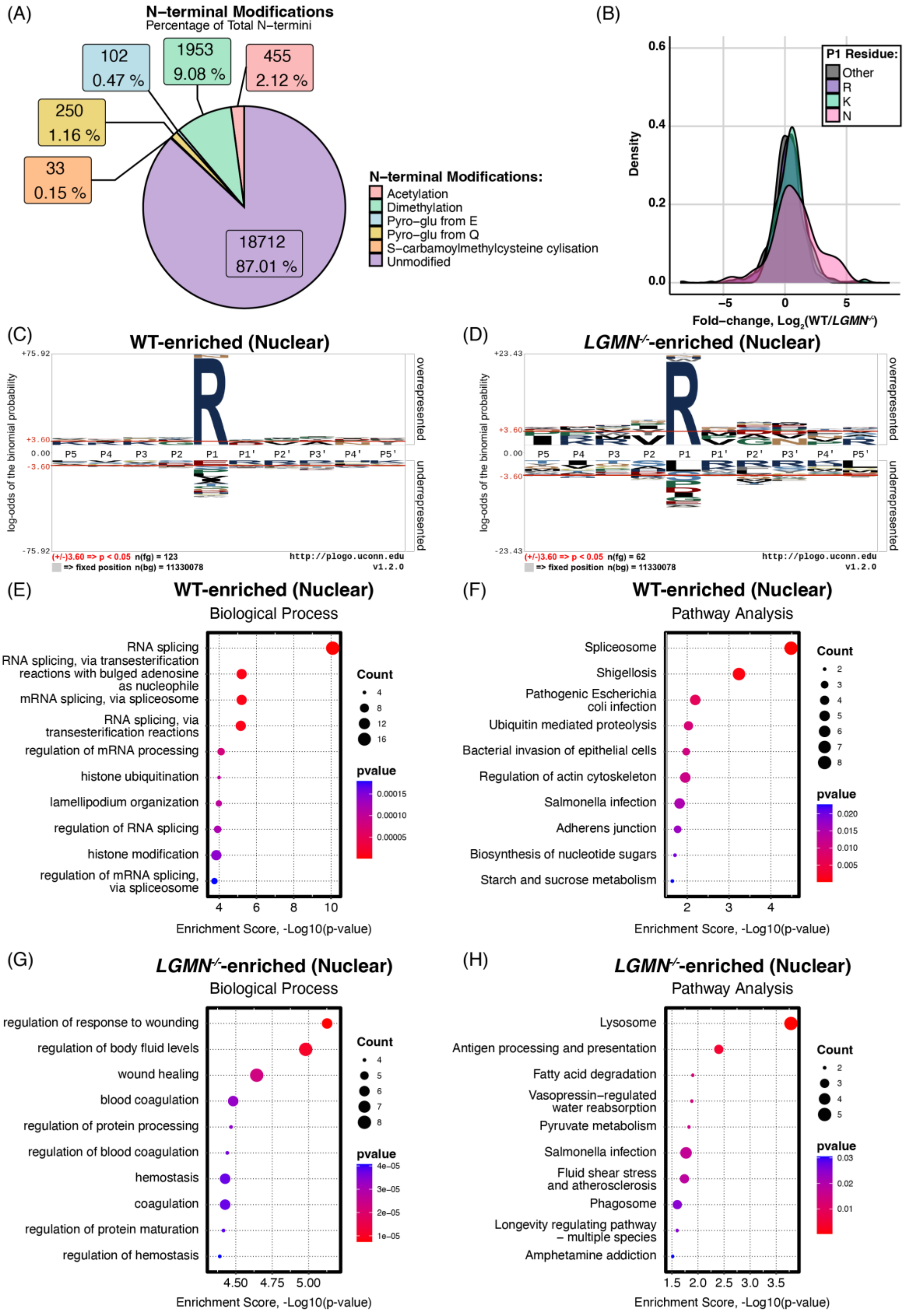


**Figure S14. Bioinformatic analysis of wild-type (WT) and legumain-deficient (*LGMN*$^{-/-}$)-enriched N-termini reveals roles for nuclear cathepsin L in the spliceosome.** (A) Proportion of N-terminal modifications of the total peptide identified by mass spectrometry analysis in the No-enrichment Identification of Cleavage Events (NICE) workflow across all biological replicates (n = 4 per group). (B-C) Sequence logo for the identified cleavage sites in the WT-enriched (B) and *LGMN*$^{-/-}$-enriched (C) N-termini. Figures were created using pLogo with significance indicated by the red line ($p < 0.05$). (D-G) Gene ontology and pathway analysis of WT-enriched (D-E) and *LGMN*$^{-/-}$-enriched (F-G) N-termini using SRplot.

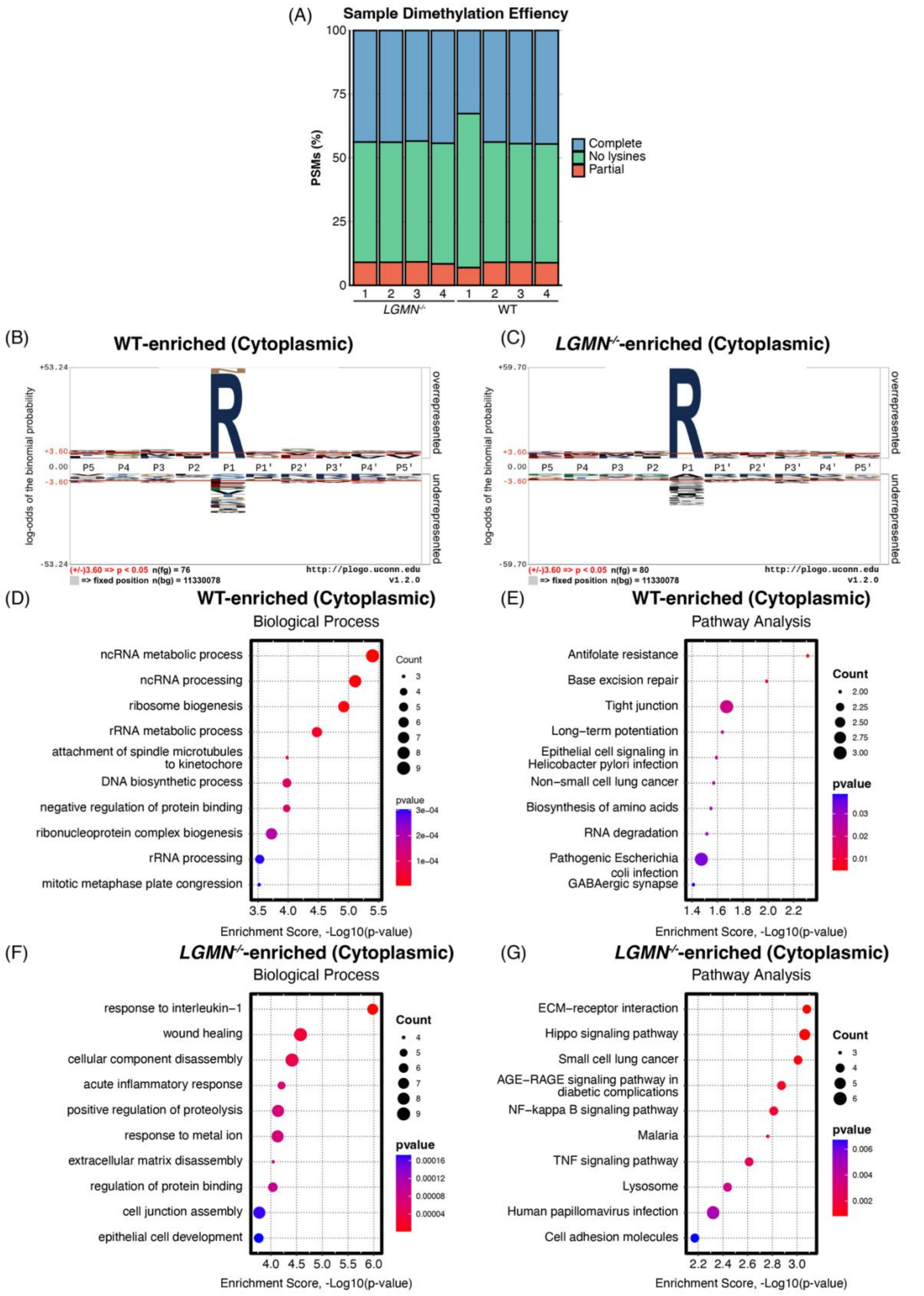


**Figure S15. Bioinformatic analysis of wild-type (WT) and legumain-deficient (*LGMN*$^{-/-}$)-enriched N-termini from cytoplasmic-enriched fractions suggests roles in RNA processing.** (A) Dimethylation labelling efficiency in cytoplasmic-enriched fractions collected from WT and *LGMN*$^{-/-}$ HSC-3 cells (n = 4 per group). Peptides with all internal lysines dimethylated represent complete dimethylation (blue), whilst those with lysines missing dimethyl labelling are incomplete (red). Peptides with no internal lysines are also shown (green). (B-C) Sequence logo for the identified cleavage sites in the WT-enriched (B) and *LGMN*$^{-/-}$-enriched (C) N-termini. Figures were created using pLogo with significance indicated by the red line ($p < 0.05$). (D-G) Gene ontology and pathway analysis of WT-enriched (D-E) and *LGMN*$^{-/-}$-enriched (F-G) N-termini using SRplot.
