## Supplementary material for "Legumain drives processing of cathepsins and nuclear localisation of cathepsin L": Table S1

**Table S1. CRISPR-Cas9 single cell clone sequencing of knockouts.**

<table>
<tr><th>Clone<br>Knockout Score (Indel %)[1]</th><th>Indel</th><th>Contribution</th><th>Sequence</th></tr>
<tr><td>LGMN WT<br>HSC-3<br>(Control)</td><td>-</td><td>-</td><td>CGTACATCATCACAACGATCTGTTCGT<br>CAGGAATCCCATTGCGGTGAATGATCTGGTAGGCATGGCACGC<br>GTCTG</td></tr>
<tr><td rowspan="7">LGMN$^{-/-}$ KO<br>HSC-3<br>9 (90%)</td><td>-18</td><td>33%</td><td>CGTACAT------------------<br>|GTCAGGAATCCCATTGCGGTGAATGATCTGGTAGGCATGGCAC<br>GCGTCTG</td></tr>
<tr><td>-3</td><td>20%</td><td>CGTACATCATCACAACGATCTGTTC|---<br>AGGAATCCCATTGCGGTGAATGATCTGGTAGGCATGGCACGCG<br>TCTG</td></tr>
<tr><td>-18</td><td>13%</td><td>CGTACATCATCA-------------|-----<br>GAATCCCATTGCGGTGAATGATCTGGTAGGCATGGCACGCGTCT<br>G</td></tr>
<tr><td>-3</td><td>9%</td><td>CGTACATCATCACAACGATCTG---<br>|GTCAGGAATCCCATTGCGGTGAATGATCTGGTAGGCATGGCAC<br>GCGTCTG</td></tr>
<tr><td>-24</td><td>8%</td><td>CGTACAT------------------|------<br>AATCCCATTGCGGTGAATGATCTGGTAGGCATGGCACGCGTCTG</td></tr>
<tr><td>-3</td><td>6%</td><td>CGTACATCATCACAACGATCTGTT-|--<br>CAGGAATCCCATTGCGGTGAATGATCTGGTAGGCATGGCACGC<br>GTCTG</td></tr>
<tr><td>-24</td><td>1%</td><td>CGTACATC-----------------|-------<br>ATCCCATTGCGGTGAATGATCTGGTAGGCATGGCACGCGTCTG</td></tr>
<tr><td>LGMN WT<br>SCC-9<br>(Control)</td><td>-</td><td>-</td><td>CGTACATCATCACAACGATCTGTTCGT<br>CAGGAATCCCATTGCGGTGAATGATCTGGTAGGCATGGCACGC<br>GTCTG</td></tr>
<tr><td>LGMN$^{-/-}$ KO<br>SCC-9<br>99 (99%)</td><td>-29</td><td>99%</td><td>CGTACATCAT---------------|--------------<br>TGCGGTGAATGATCTGGTAGGCATGGCACGCGTCTG</td></tr>
<tr><td>LGMN WT<br>RAW264.7<br>(Control)</td><td>-</td><td>-</td><td>CCACCTGCCCGACGACATCAACGGTAGGTATCAAACAGGCTGA<br>GGTTGGTTGTCATTGGATATTGCCCGTTTAATACCAAGCGTTCCA<br>CCTGTGCAGGGCCACTCAGGGCTGCTTGCTATCTAGGTGGAGG<br>TCATGCCCCCTGTGGTCGGGTCTGGTTTCTACTCCACAAAA</td></tr>
<tr><td rowspan="4">LGMN$^{-/-}$ KO<br>RAW264.7<br>Replicate 1<br>49 (93%)</td><td>-11</td><td>49%</td><td>CCACCTGCCCGACG-----------|--<br>GTATCAAACAGGCTGAGGTTGGTTGTCATTGGATATTGCCCGTTT<br>AATACCAAGCGTTCCACCTGTGCAGGGCCACTCAGGGCTGCTT<br>GCTATCTAGGTGGAGGTCATGCCCCCTGTGGTCGGGTCTGGTTT<br>CTACTCCACAAAA</td></tr>
<tr><td>-9</td><td>23%</td><td>CCACCTGCCCGACGACA--------|-<br>GGTATCAAACAGGCTGAGGTTGGT<br>TGTCATTGGATATTGCCCGTTTAATA<br>CCAAGCGTTCCACCTGTGCAGGG<br>CCACTCAGGGCTGCTTGCTATCTA<br>GGTGGAGGTCATGCCCCCTGTGG<br>TCGGGTCTGGTTTCTACTCCACAAAA</td></tr>
<tr><td>-9</td><td>11%</td><td>CCACCTGCCCGACGACATC------|---<br>TATCAAACAGGCTGAGGTTGGTTGTCA<br>TTGGATATTGCCCGTTTAATACCAAGC<br>GTTCCACCTGTGCAGGGCCACTCAGG<br>GCTGCTTGCTATCTAGGTGGAGGTCAT<br>GCCCCCTGTGGTCGGGTCTGGTTTCT<br>ACTCCACAAAA</td></tr>
<tr><td>-9</td><td>10%</td><td>CCACCTGCCCGACGACAT-------|--<br>GTATCAAACAGGCTGAGGTTGGTTG<br>TCATTGGATATTGCCCGTTTAATACC<br>AAGCGTTCCACCTGTGCAGGGCCA<br>CTCAGGGCTGCTTGCTATCTAGGTG<br>GAGGTCATGCCCCCTGTGGTCGGG<br>TCTGGTTTCTACTCCACAAAA</td></tr>
</table>

| | | | |
|---|---|---|---|
| *LGMN*$^{-/-}$ KO RAW264.7 Replicate 2 49 (92%) | -13 | 49% | CCACCTGCCCGACG-----------\|--GTATCAAACAGGCTGAGGTTGGTTGTCATTGGATATTGCCCGTTTAATACCAAGCGTTCCACCTGTGCAGGGCCACTCAGGGCTGCTTGCTATCTAGGTGGAGGTCATGCCCCCTGTGGTCGGGTCTGGTTTCTACTCCACAAAA |
| | -9 | 23% | CCACCTGCCCGACGACA--------\|-GGTATCAAACAGGCTGAGGTTGGTTGTCATTGGATATTGCCCGTTTAATACCAAGCGTTCCACCTGTGCAGGGCCACTCAGGGCTGCTTGCTATCTAGGTGGAGGTCATGCCCCCTGTGGTCGGGTCTGGTTTCTACTCCACAAAA |
| | -9 | 11% | CCACCTGCCCGACGACATC------\|---TATCAAACAGGCTGAGGTTGGTTGTCATTGGATATTGCCCGTTTAATACCAAGCGTTCCACCTGTGCAGGGCCACTCAGGGCTGCTTGCTATCTAGGTGGAGGTCATGCCCCCTGTGGTCGGGTCTGGTTTCTACTCCACAAAA |
| | -9 | 9% | CCACCTGCCCGACGACAT-------\|--GTATCAAACAGGCTGAGGTTGGTTGTCATTGGATATTGCCCGTTTAATACCAAGCGTTCCACCTGTGCAGGGCCACTCAGGGCTGCTTGCTATCTAGGTGGAGGTCATGCCCCCTGTGGTCGGGTCTGGTTTCTACTCCACAAAA |
| *LGMN*$^{-/-}$ KO RAW264.7 Replicate 3 48 (93%) | -13 | 48% | CCACCTGCCCGACG-----------\|--GTATCAAACAGGCTGAGGTTGGTTGTCATTGGATATTGCCCGTTTAATACCAAGCGTTCCACCTGTGCAGGGCCACTCAGGGCTGCTTGCTATCTAGGTGGAGGTCATGCCCCCTGTGGTCGGGTCTGGTTTCTACTCCACAAAA |
| | -9 | 23% | CCACCTGCCCGACGACA--------\|-GGTATCAAACAGGCTGAGGTTGGTTGTCATTGGATATTGCCCGTTTAATACCAAGCGTTCCACCTGTGCAGGGCCACTCAGGGCTGCTTGCTATCTAGGTGGAGGTCATGCCCCCTGTGGTCGGGTCTGGTTTCTACTCCACAAAA |
| | -9 | 15% | CCACCTGCCCGACGACATCAA----\|-----TCAAACAGGCTGAGGTTGGTTGTCATTGGATATTGCCCGTTTAATACCAAGCGTTCCACCTGTGCAGGGCCACTCAGGGCTGCTTGCTATCTAGGTGGAGGTCATGCCCCCTGTGGTCGGGTCTGGTTTCTACTCCACAAAA |
| | -9 | 5% | CCACCTGCCCGACGACAT-------\|--GTATCAAACAGGCTGAGGTTGGTTGTCATTGGATATTGCCCGTTTAATACCAAGCGTTCCACCTGTGCAGGGCCACTCAGGGCTGCTTGCTATCTAGGTGGAGGTCATGCCCCCTGTGGTCGGGTCTGGTTTCTACTCCACAAAA |
| | -9 | 2% | CCACCTGCCCGACGACATCA-----\|----ATCAAACAGGCTGAGGTTGGTTGTCATTGGATATTGCCCGTTTAATACCAAGCGTTCCACCTGTGCAGGGCCACTCAGGGCTGCTTGCTATCTAGGTGGAGGTCATGCCCCCTGTGGTCGGGTCTGGTTTCTACTCCACAAAA |
| *CTSL* WT HSC-3 (Control) | - | - | ACCGGCTTTGTGGACATCCCTAAGCAGGAGAAGGCCCTGATGAAGGCAGTTGCAACTGTGGGGCCCATTTCTGTTGCTATTGATGCAGGTCATGAGTCCTTCCTGTTCTATAAAGAAGGTAAGCATATTTTTCTTTGTAGAAATTGATGCAGAAAATAGAGTATCATGAAATGAA |
| *CTSL*$^{-/-}$ KO HSC-3 Replicate 1 94 (94%) | -2 | 45% | ACCGGCTTTGTGGACATCCCTAA--\|AGGAGAAGGCCCTGATGAAGGCAGTTGCAACTGTGGGGCCCATTTCTGTTGCTATTGATGCAGGTCATGAGTCCTTCCTGTTCTATAAAGAAGGTAAGCATATTTTTCTTTGTAGAAATTGATGCAGAAAATAGAGTATCATGAAATGAA |
| | -5 | 30% | ACCGGCTTTGTGGACATCCC-----\|AGGAGAAGGCCCTGATGAAGGCAGTTGCAACTGTGGGGCCCATTTCTGTTGCTATTGATGCAGGTCATGAGTCCTTCCTGTTCTATAAAGAAGGTAAGCATATTTTTCTTTGTAGAAATTGATGCAGAAAATAGAGTATCATGAAATGAA |
| | -5 | 19% | ACCGGCTTTGTGGACATCCCTA---\|--GAGAAGGCCCTGATGAAGGCAGTTGCAACTGTGGGGCCCATTTCTGTTGCTATTGATGCAGGTCATGAGTCCTTCCTGTTCTATAAAGAAGGTAAGCATATTTTTCTTTGTAGAAATTGATGCAGAAAATAGAGTATCATGAAATGAA |
| *CTSL*$^{-/-}$ KO HSC-3 Replicate 2 | -2 | 46% | ACCGGCTTTGTGGACATCCCTAA--\|AGGAGAAGGCCCTGATGAAGGCAGTTGCAACTGTGGGGCCCATTTCTGTTGCTATTGATGCAGGTCATGAGTCCTTCCTGTTCTATAAAGAAGGTAAGCATATTTTTCTTTGTAGAAATTGATGCAGAAAATAGAGTATCATGAAATGAA |

| | | | |
|---|---|---|---|
| 94 (94%) | -5 | 32% | ACCGGCTTTGTGGACATCCC-----\|AGGAGAAGGCCCTGATGAAGGCAGTTGCAACTGTGGGGCCCATTTCTGTTGCTATTGATGCAGGTCATGAGTCCTTCCTGTTCTATAAAGAAGGTAAGCATATTTTTCTTTGTAGAAATTGATGCAGAAAATAGAGTATCATGAAATGAA |
| | -5 | 16% | ACCGGCTTTGTGGACATCCCTA---\|--GAGAAGGCCCTGATGAAGGCAGTTGCAACTGTGGGGCCCATTTCTGTTGCTATTGATGCAGGTCATGAGTCCTTCCTGTTCTATAAAGAAGGTAAGCATATTTTTCTTTGTAGAAATTGATGCAGAAAATAGAGTATCATGAAATGAA |
| *CTSL*$^{-/-}$ KO HSC-3 Replicate 3 94 (94%) | -2 | 45% | ACCGGCTTTGTGGACATCCCTAA--\|AGGAGAAGGCCCTGATGAAGGCAGTTGCAACTGTGGGGCCCATTTCTGTTGCTATTGATGCAGGTCATGAGTCCTTCCTGTTCTATAAAGAAGGTAAGCATATTTTTCTTTGTAGAAATTGATGCAGAAAATAGAGTATCATGAAATGAA |
| | -5 | 34% | ACCGGCTTTGTGGACATCCC-----\|AGGAGAAGGCCCTGATGAAGGCAGTTGCAACTGTGGGGCCCATTTCTGTTGCTATTGATGCAGGTCATGAGTCCTTCCTGTTCTATAAAGAAGGTAAGCATATTTTTCTTTGTAGAAATTGATGCAGAAAATAGAGTATCATGAAATGAA |
| | -5 | 15% | ACCGGCTTTGTGGACATCCCTA---\|--GAGAAGGCCCTGATGAAGGCAGTTGCAACTGTGGGGCCCATTTCTGTTGCTATTGATGCAGGTCATGAGTCCTTCCTGTTCTATAAAGAAGGTAAGCATATTTTTCTTTGTAGAAATTGATGCAGAAAATAGAGTATCATGAAATGAA |

[1]KO score analysed using Inference of CRISPR Edits (ICE) analysis of single cell clones, generated by https://ice.synthego.com
